# ProtoCloud: a Prototypical Self-explaining Model for Single-cell Analysis

**DOI:** 10.64898/2026.02.06.704364

**Authors:** Kaiyun Guo, Jiarui Ding

**Affiliations:** Department of Computer Science, University of British Columbia, Vancouver, British Columbia, Canada, V6T 1Z4

**Keywords:** single-cell RNA-sequencing, cell state, rare cell type, prototypical network, self-explaining model, disentanglement, layer-wise relevance propagation, prototypical relevance propagation, deep generative models, variational autoencoder

## Abstract

Cell type annotation is a fundamental task in single-cell genomics. Although various methods have been developed for automatic cell type annotation, they often function as black-box models, making predictions without explaining their reasoning and lacking proper uncertainty estimation for their predictions. Furthermore, they often struggle to annotate rare cell types. We introduce ProtoCloud, a self-explaining deep generative model trained end-to-end to embed cells into a structured, low-dimensional space organized around cell type-specific prototypes. Coupled with a specifically-designed data augmentation strategy, it matches or outperforms existing methods in cell type annotation across 11 large-scale datasets, particularly for rare cell types. Moreover, ProtoCloud improves data annotation quality by identifying and re-annotating mis-annotated training cells through a built-in certainty quantification mechanism based on cell-prototype similarity. Finally, ProtoCloud provides interpretable predictions by identifying key genes that drive its classifications, facilitating the discovery of both known and novel cell type marker genes. Applied to a time-course dataset of post-injury retinal neurons, ProtoCloud successfully annotates previously unassigned cells; on the esophageal cell atlas, it identifies rare but potentially important cell populations and their marker genes relevant to esophageal inflammation.

## 1 Introduction

In recent years, technological advancements in single-cell genomics, especially single-cell and single-nucleus RNA sequencing (scRNA-seq and snRNA-seq)^1–7^, have enabled the construction of comprehensive tissue atlases^8–11^, each encompassing hundreds of thousands to millions of cells. These atlases provide detailed insights into tissue cellular composition and intercellular interactions^12–14^, cell type-specific and shared biological processes^15–17^, in development^18–20^, homeostasis^21^, aging^22^, and inflammation and disease^12,23–25^. Computational methods have been developed to map newly sequenced cells to these reference atlases, facilitating rapid cell type annotation and accelerating biological discoveries^26–29^. This supervised, reference mapping approach contrasts with unsupervised clustering strategies, which rely on either manual annotation^30^ or automated marker gene scoring^31^ for cell type assignment, and remain indispensable in settings where annotated reference datasets are unavailable.

Among the reference-mapping approach, deep learning models have emerged as powerful tools for analyzing single-cell datasets^26,32–41^. Because of their scalability in processing large-scale datasets, flexibility in handling different data modalities with potential experimental-specific nuisance factors, and effectiveness in analyzing complex high-dimensional data, deep learning models are well-suited for modeling scRNA-seq datasets. Despite these successes, these models often function as black boxes—excelling at predictions but offering limited interpretability regarding their decision-making process^42^. For large-scale cell type annotation, an ideal deep learning model should be self-explaining, offering not only accurate predictions but also clear insights into its decision-making process^43^. For example, if a cell is predicted to be a B cell, a self-explaining model should provide dual explainability: 1) gene-level features highlighting key expressed genes driving the decision, and 2) cell-level similarities to reference B cell populations or B cell prototypes. Such transparency is crucial for assessing prediction uncertainties and building reliable cell atlases.

Single-cell data present additional challenges that complicate accurate cell type annotation, even with deep learning methods. First, tissues typically consist of dominant and rare cell types, complicating the identification of rare cell populations^25,44^. The origins of these rare cell types may be biological or technical. For instance, basophils play essential roles in allergy, but they account for less than 1% of circulating blood cells^45^. Conversely, eosinophils are enriched in the diseased human esophagus, yet these cells are frequently undetected in single-cell studies due to technical limitations in conventional scRNA-seq pipelines^46–48^. Second, cell type-irrelevant nuisance factors introduce another layer of complexity, further hindering accurate cell type annotation and interpretation^49,50^. While specialized tools have been developed to mitigate these nuisance factors^33,51^, identifying which specific factors require correction in a given dataset remains challenging.

Motivated by these gaps, we introduce ProtoCloud, a self-explaining deep learning model that achieves state-of-the-art performance in single-cell cell type annotation with both cell-level and gene-level explainability. The model embeds cells around cell type-specific prototypes in a low-dimensional latent space, quantifying prediction uncertainty through distance-based similarities between cell embeddings and the closest prototypes of the predicted cell types. By backpropagating cell-prototype similarities in the latent space to the observed gene space, it highlights the most relevant genes that drive the model’s decisions. Furthermore, ProtoCloud partitions its latent space into two components that isolate cell type information from cell type-irrelevant factors^52,53^. To improve performance on rare cell types, the model employs a unique data augmentation technique that models gene counts using multinomial distribution-based sampling. Through its model architecture, loss functions, and training strategy, ProtoCloud effectively captures the structures of complex high-dimensional scRNA-seq data in the latent space, resulting in accurate and robust predictions with meaningful biological insights.

Across multiple datasets, ProtoCloud performs favorably compared to several widely used scRNA-seq annotation tools. It also successfully disentangles cell types from technical batches in the latent embeddings. We further demonstrate its effectiveness by reannotating potentially mis-annotated cells in a widely used peripheral blood mononuclear cell dataset^3,33^. In a time-course study of retinal ganglion cells following acute injury^54^, ProtoCloud successfully annotates previously unassigned cells and outperforms competing methods. Additionally, when trained on an esophageal mucosal cell atlas^25^ (421,312 cells, 72 cell types) to annotate cells from two additional studies, ProtoCloud identifies disease-associated rare cell populations, including the novel *PRDM16* ^+^ dendritic cells in both test cohorts. Collectively, these results highlight ProtoCloud’s versatility in refining reference cell type annotations, identifying rare cell populations, and supporting the construction of comprehensive cell atlases, establishing it as a valuable tool for single-cell data analysis.

## 2 RESULTS

ProtoCloud is built upon a variational autoencoder (VAE) architecture trained end-to-end for single-cell genomics analysis^55,56^. Requiring only cell UMI counts and reference cell labels as inputs during training, it employs a probabilistic encoder to project high-dimensional single-cell data into a low-dimensional latent space and a probabilistic decoder to reconstruct the data from this latent space back to the original gene expression space (**Figure 1**A). To handle complex single-cell data, which contains both cell type-specific signals and various technical artifacts, ProtoCloud employs a carefully designed structured latent space. The model partitions its latent space into two disjoint subspaces, each dedicated to capturing a different functional aspect of the data.

**Figure 1:**
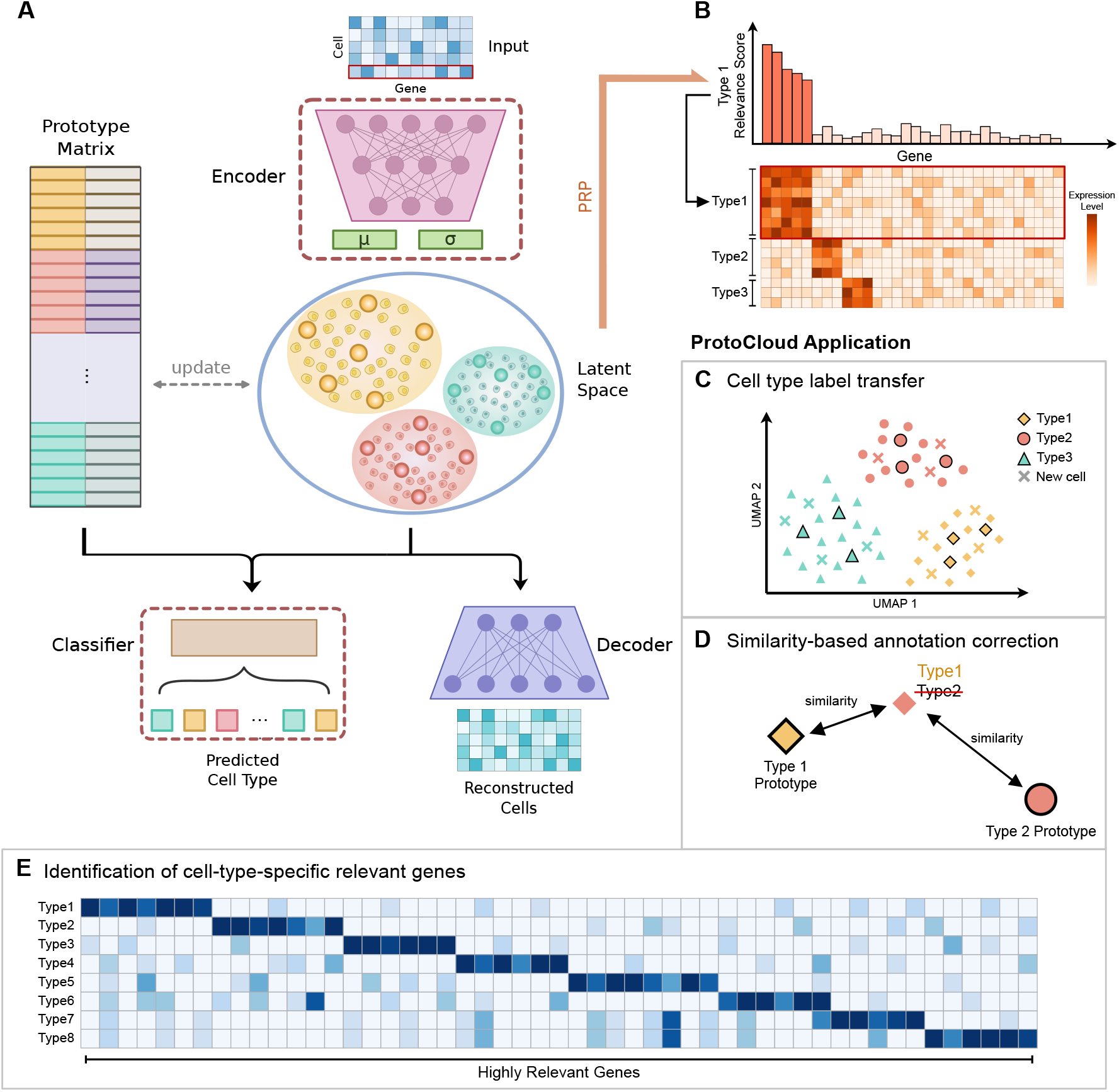
ProtoCloud overview. **(A)** An illustration of the ProtoCloud model. The model has four major components: a probabilistic encoder, a probabilistic decoder, a prototype matrix, and a linear classifier. The model takes only raw UMI counts (cell-by-gene count matrix) as input during inference. Cell inputs are encoded into a low-dimensional latent space (*d*_*z*_ = 20 by default), with different colors indicating cell types. Larger points in the latent space represent prototypes, which are pre-initialized (six per cell type) and share the same latent space as the cell embeddings. Cell type information (brighter color) is encoded in the first half of the latent dimensions. The latent embeddings are used for both cell type prediction, based on similarity to the prototypes, and for reconstructing the gene expression through the decoder. **(B)** ProtoCloud provides inherent interpretability through prototypical relevance propagation (PRP), generated simultaneously once the model is trained. Each prototype undergoes PRP to produce gene-level relevance scores. Genes with higher relevance scores are the decision-relevant genes of the corresponding cell type. **(C – E)** Applications of ProtoCloud. **(C)** The model enables accurate and robust transfer of cell type annotations across datasets. Shapes denote ground truth cell types, and colors indicate predicted labels. Symbols with black edges represent prototypes, while those without edges are cell embeddings. Crosses mark newly added query cells predicted to the corresponding cell types. **(D)** ProtoCloud’s similarity-based classification process enables detection of anomalous annotations, improving the quality of the ground-truth annotations. A cell with ground truth Type 1 (diamond shape) was initially mislabeled as Type 2. ProtoCloud corrects this annotation by assigning the cell to Type 1 based on a higher similarity score. **(E)** The identified cell type-relevant genes can act as candidate markers and provide molecular insights into cell identities.

The first half of the latent embedding is designed to capture cell type information, whereas the second half is intended to capture other factors, such as batch effects (**Methods**). This strategic division allows ProtoCloud to perform cell type predictions and generate explanations based on the cell type-relevant portion of the latent space, while still using the complete latent representation to accurately reconstruct cells’ gene expression profiles.

Each cell type is associated with a set of prototypes that also reside in the latent space. These prototypes are designed to capture: (1) inter-cell type variation, ensuring that prototypes of the same cell type are proximal to each other while those of different types are well-separated, and (2) intra-cell type variation, enabling the prototypes of a given cell type to capture its inherent heterogeneity. For cell type annotation, ProtoCloud achieves interpretability at two levels. To achieve cell level interpretability of a prediction, cell type classification is based on the similarities of a cell’s embedding to these prototypes (**Methods**). For gene-level interpretability of a prediction, ProtoCloud applies prototypical relevance propagation (PRP)^57–59^ to the trained prototypes of the predicted class to identify decision-driving genes. For each cell, it assigns a relevance score to each gene by backpropagating the initial relevance, which is positive for the prototypes of the cell to which it is assigned and negative for other prototypes, through the encoder network to the input space (**Figure 1**B and **Methods**).

By using consistent hyperparameters across all datasets, we demonstrate the robustness and versatility of ProtoCloud across diverse biological contexts. We highlight three major applications of ProtoCloud in scRNA-seq data analysis: (1) reliable label transfer between datasets (**Figure 1**C), (2) detection and correction of annotation anomalies to improve data quality (**Figure 1**D), and (3) identification of candidate cell type marker genes (**Figure 1**E).

### Accurate and robust cell type annotation

To evaluate ProtoCloud’s performance in cell type annotation, we performed a comparative analysis against eight state-of-the-art methods, including two probabilistic machine learning approaches (Seurat^27^ and CellTypist^28^), four deep learning models (scANVI^26^, TOSICA^60^, scPoli^61^, and SIMS^62^), and two NLP-inspired foundational models (scGPT^40^ and scBERT^38^). We benchmarked these methods across eight diverse datasets spanning multiple species, organs, and sequencing technologies^3,8,25,54,63,64^ (**Methods**). Model performance was assessed across multiple random seeds, with average accuracy, macro F1 score, and Cohen’s kappa coefficient^65^ reported as summary metrics (**Methods**). Using default hyperparameters across all datasets, ProtoCloud exhibited strong and consistent performance, often matching or surpassing the baselines across datasets and evaluation metrics (**Figures 2**A– **2**C, **Table** S1, **Methods**). Other state-of-the-art methods such as scANVI also showed high accuracy, but requires batch information as input. Importantly, ProtoCloud demonstrated notably higher accuracy for low-abundance cell types, as reflected in the macro F1 score (**Figure 2**D, **Table** S2). These results underscore ProtoCloud’s ability to achieve robust and reliable classification performance, even for rare cell types. As a deep learning model trained with mini-batch, ProtoCloud is scalable to process large datasets (**Figure** S1A, **Methods**).

**Figure 2:**
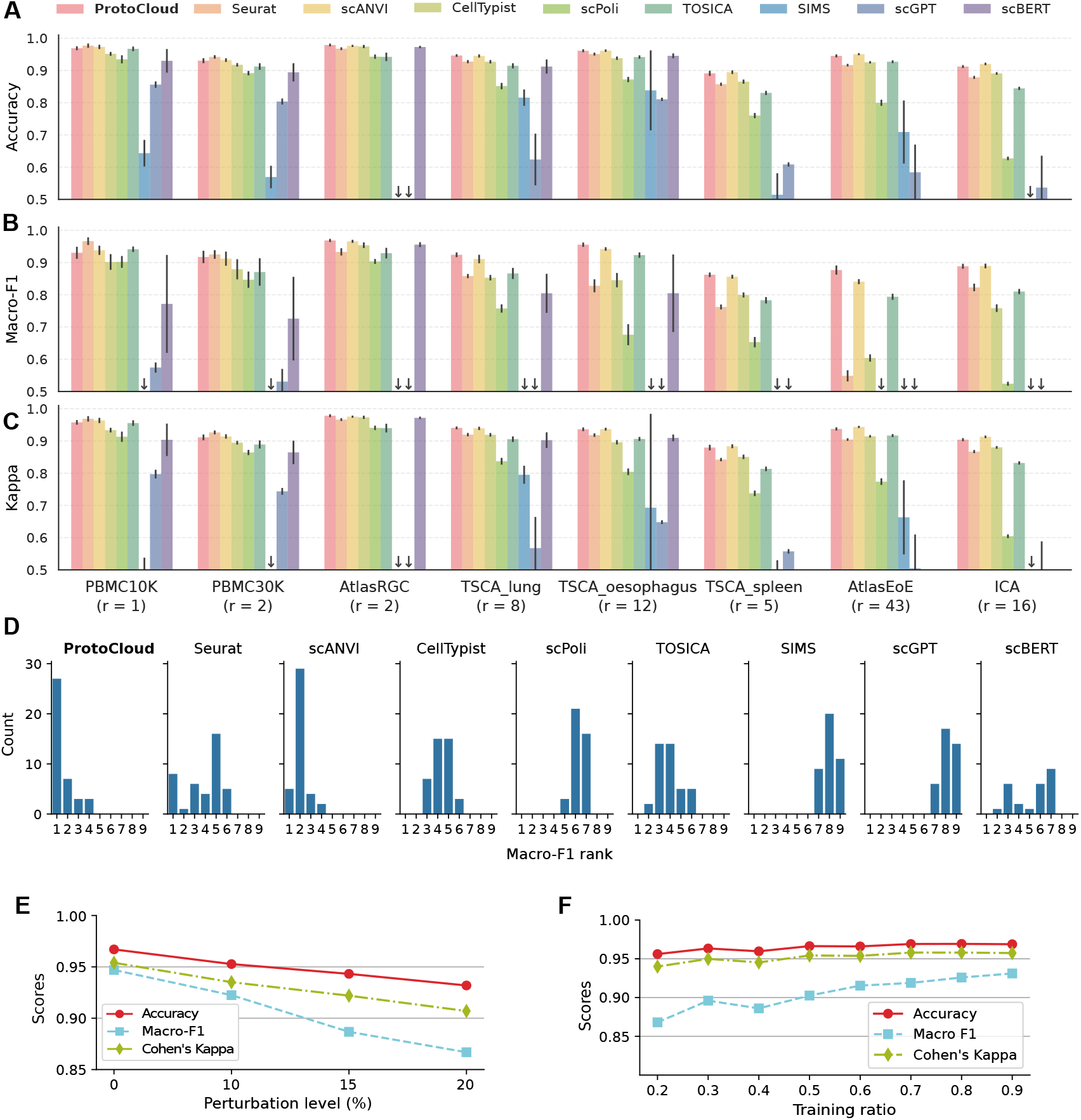
Performance of ProtoCloud on cell type classification. **(A – C)** Benchmarking of cell type annotation performance. We compared ProtoCloud against Seurat V4^27^, scANVI^26^, CellTypist^28^, scPoli^61^, TOSICA^60^, SIMS^62^, scGPT^40^, and scBERT^38^ for cell type annotation across eight datasets, ranging from approximately 10,000 to 400,000 cells. The x-axis represents datasets, with the numbers in the parentheses indicating the number of rare cell types in each dataset. Downward-pointing arrows indicate values below 0.5. Error bars: standard error of the mean. **(A)** Evaluation metrics include accuracy, **(B)** macro F1 score, **(C)** and Cohen’s kappa coefficient. The metrics were averaged over five random seed experiments, with 80% of the data used for training and 20% for validation in each run. **(D)** Macro F1 score rank distributions across methods and datasets over all experimental repetitions. **(E, F)** Model performance analysis under varying conditions using the PBMC10K dataset^3^. **(E)** Validation accuracy as the proportion of label perturbation in the training set increases from 0% to 20%. **(F)** Validation accuracy as training data ratios decrease, ranging from 0.8 to 0.1.

To evaluate ProtoCloud’s robustness, we assessed its performance under varying conditions using the peripheral blood mononuclear cell (PBMC) dataset^3,33^ We introduced perturbations by randomly shuffling different proportions of the training labels (i.e., 10%, 15%, and 20%). Even under these label perturbations, the model maintained high validation accuracies above 93.18% (**Figure 2**E), demonstrating its resilience to label noise. We further investigated ProtoCloud’s performance under limited training data scenarios by varying the training data ratio from 0.10 to 0.80. The model maintained stable performance across these training set sizes, with accuracy ranging from 95.34% to 97.64% (**Figure 2**F). Notably, even when trained on only 10% of the data, the macro F1 score remained high at 0.87, indicating robust performance even with limited training data. The consistently strong performance and robustness demonstrated by ProtoCloud in cell type classification tasks underscore its reliability in scRNA-seq analysis and form the basis for its robust explainability.

### Batch-separated informative latent space

Despite not requiring batch information as input, ProtoCloud’s two-stage curriculum and structured latent space design (**Methods**) effectively separate and capture batch-related variability in its latent subspace. To assess this separation in the latent space, we used two datasets with known batch annotations: PBMC30K^63^ and AtlasRGC^54^.

We assessed the information content of the separated latent spaces by calculating biological conservation and batch mixing metrics (**Figure 3**A, **Methods**). Consistent with our design, the first component, **z**^1^, is highly enriched for biological information, evidenced by a near-perfect cell type local inverse Simpson’s Index (cLISI) of 1.00 and high clustering evaluation metrics (e.g., the normalized mutual information metric NMI = 0.95). Conversely, the second half of the latent embedding (**z**^2^) shows substantially lower biological conservation (e.g., cLISI = 0.78 and NMI = 0.06) but preserves batch effects, with an integration local inverse Simpson’s Index (iLISI) of zero, compared to 0.44 from **z**^1^. These trends are consistent across additional evaluation metrics (e.g., KBET and Graph connectivity), with the exception of the silhouette batch metric, which is known to be unreliable for assessing batch correction^66^.

**Figure 3:**
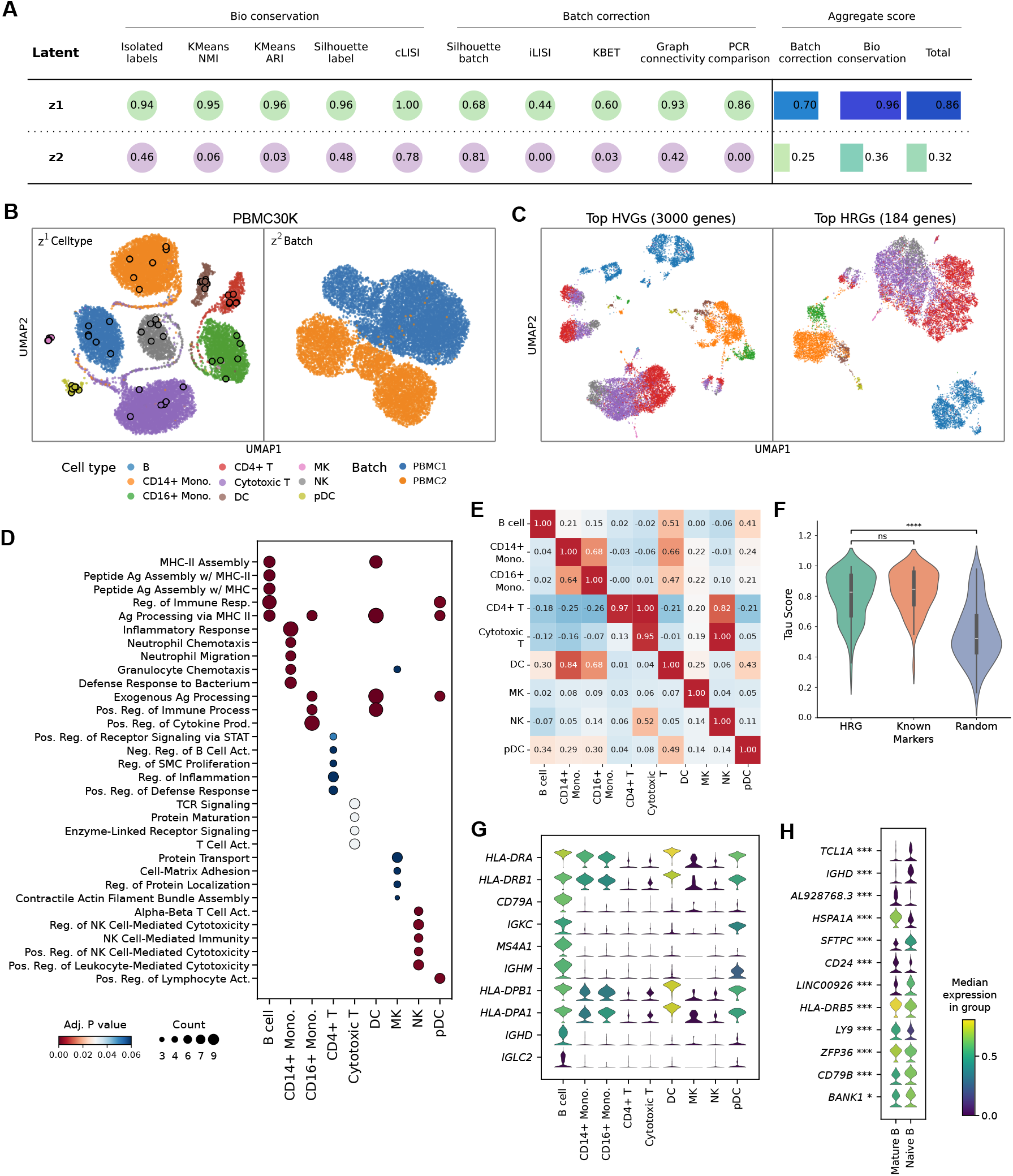
Latent space organization and HRG verification in ProtoCloud. **(A)** Evaluation of latent space disentanglement using the PBMC30K^63^ dataset. Comparison of scIB metrics for the biological subspace (first-half latent space, **z**^1^) and the batch subspace (second-half latent space, **z**^2^). Metrics are grouped into biological conservation and batch correction categories. A score closer to 1 indicates better performance. **(B)** Visualization of ProtoCloud latent representations of PBMC30K. UMAP projections show cell type-specific clustering in **z**^1^ (left) and batch effects in **z**^2^ (right) of the latent space. Prototypes are represented by dots with black outlines. **(C)** Visualization of ProtoCloud latent representations of PBMC30K. UMAP projections show the representation using all 3,000 HVGs (left) compared to the representation using the union of the top 30 HRGs of each cell type (right), resulting in 184 unique genes in total. **(D)** Gene ontology (GO) biological process enrichment analysis of HRGs across cell types in PBMC30K. The dot plot displays the top enriched pathways for each cell type. Dot size represents the number of HRGs associated with the pathway, and color indicates statistical significance. **(E)** Cell type specificity of HRGs in PBMC30K. Heatmaps comparing row-normalized gene signature scores based on the top 15 HRGs across nine cell types. Diagonal elements (matching cell types) represent on-target specificity, where high values indicate relatively high expression in the respective cell type, while off-diagonal elements represent off-target expression. **(F)** Quantitative comparison of gene specificity across three gene sets in PBMC30K. Violin plots display the distributions of Tau specificity scores for HRGs, canonical marker genes, and randomly selected genes. HRGs exhibit cell type specificity comparable to known markers, achieving a median Tau score of 0.828, while randomly selected genes show significantly lower specificity. **(G)** Expression profiles of the top ten B cell-specific HRGs in PBMC30K. The y-axis lists the top ten B cell-specific HRGs in descending rank of relevance. Expression distributions of these top ten B cell-specific HRGs across cell types are shown. High expression in B cells (leftmost column) contrasted with low expression in other cell types demonstrates strong cell type specificity. **(H)** Expression profiles of the top differentially ranked HRGs between mature and naïve B cells in the TSCA lung^64^ dataset. Differential expression significance was assessed using the Wilcoxon rank-sum test with Benjamini-Hochberg correction, denoted by asterisks (* *p* ≤ 0.05, *** *p* ≤ 0.001).

To visualize the separation of information encoded in different segments of the latent space, we used Uniform Manifold Approximation and Projection (UMAP)^67^ to project both the first and the second halves of the latent space into two dimensions. Similarly, the UMAP of the first half of the latent space (**z**^1^) shows clear separation by cell types (**Figure 3**B), but no apparent clustering by batch (**Figure** S1B), whereas the UMAP of the second half of the latent space (**z**^2^) reveals predominant grouping of cells by batch, with mixed cell type composition within these batch groups. Direct visualization of the latent space reveals that batch effects are confined to specific dimensions of the latent space (**Figure** S1C). The consistent pattern is also observed in the AtlasRGC dataset, with cells grouped by type in the first half of the latent space and by batch in the second half of the latent space (**Figures** S1D–S1F). This clear dimensional disentanglement suggests that different components of the latent space effectively encode distinct technical (batch) and biological (cell type) sources of variation.

We performed ablation studies to evaluate the impact of different training strategies on latent space organization (**Methods**). On the UMAP of the latent representation obtained from the model with a non-separated latent space, marked batch effects were observed with a low Batch Entropy Score (BES) of 0.014 (**Figure** S2A, **Methods**). We then evaluated the impact of our two-stage curriculum training strategy compared to a single-stage end-to-end approach. While both strategies maintained high biological conservation in **z**^1^, the two-stage approach demonstrated superior disentanglement capabilities (**Figure** S2B). Specifically, the two-stage training yielded a significantly higher aggregate batch correction score of 0.78 for **z**^1^, compared to 0.67 for the single-stage model. Next, we dissected the contribution of each loss component (**Figure** S2C). The baseline VAE configuration (without the atomic and orthogonal losses; **Methods**) resulted in collapsed prototypes with intra-class similarities approaching 1.0, indicating insufficient within-class diversity. Adding only the atomic loss improved cell-type separation (decreased inter-class similarity), although prototypes remained overly compact (high intra-class similarity). Incorporating only the orthogonal loss improved (reduced) intra-class similarity to 0.953 but increased inter-class similarity to 0.218, compared to 0.130 in the standard configuration. These results confirm that the complete loss formulation in our standard setting learns prototypes that better capture both between-cell type separation and within-cell type diversity (**Figure** S2D). Together, these ablation studies demonstrate that each auxiliary component contributes to ProtoCloud’s overall performance.

### Gene-level explainability of cell predictions

ProtoCloud employs prototypical relevance propagation (PRP, **Methods**)^57–59^ to identify the key genes driving its classification decisions. This approach provides a direct, data-driven justification for the model’s predictions. Genes assigned high relevance scores are referred to as highly relevant genes (HRGs). Users can leverage HRGs for downstream analyses instead of relying only on prior knowledge of established cell-type markers^68–70^, ensuring broad applicability across diverse biological contexts (**Figure** S3A).

To validate the representational power of HRGs, we visualized the UMAP using only the union of the top 30 HRGs of each type and compared it against the baseline constructed from top 3,000 HVGs (**Figure 3**C). Despite utilizing approximately 16-fold fewer genes, the HRG-based embedding successfully preserved the global topology of the data (e.g., the continuum arrangement of CD4^+^ T cells to Cytotoxic T cells, and then to NK cells in the UMAP; similarly, CD14^+^ monocytes and CD16^+^ monocytes are close to each other). We next assessed the biological validity of these signatures through Gene Ontology (GO) enrichment analysis (**Figure 3**D). The enriched pathways exhibited high lineage specificity, strictly aligning with known cellular functions. For instance, CD14^+^ monocytes were enriched for inflammatory response and defense response to bacterium, consistent with their primary phagocytic role. In contrast, CD16^+^ monocytes were distinctively characterized by immune regulation and antigen presentation pathways. These findings align with the established roles of classical monocytes as first responders to bacterial infection and non-classical monocytes as patrolling immune modulators.

To quantify the cell-type specificity of the selected HRGs, we calculated the gene scores^71^ using the top 15 HRGs of each cell type (**Figure 3**E). The resulting correlation matrix revealed a sharp diagonal structure, indicating that HRGs are mainly relevant to their respective cell identities, with little off-target assignment. We systematically compared ProtoCloud HRGs against curated lists of known cell-type markers from scType^31^. The top cell type-specific HRGs also overlap with differentially expressed genes specific to cell types (**Figure** S3B). As expected, the specificity index (Tau score^72^) of HRGs (*τ* = 0.801) was significantly higher than that of random background genes, and comparable to the curated markers (**Figure 3**F). Furthermore, HRGs showed elevated expression levels in their own cell types and distinctive expression profiles across other populations (**Figure 3**G, **Figure** S3C).

The relevance of HRGs highlighted subtle biological differences between closely related cell subtypes. For example, differential HRGs between mature and naïve B cells identified key distinguishing genes (**Figure 3**H, **Methods**). Highly relevant genes such as *CD79B, BANK1*, and MHC class II genes were more highly expressed in naïve B cells, while *LY9* was more specific to mature B cells.

We further investigated the consistency of HRGs across different organs using three datasets (lung, esophagus, and spleen) from the Tissue Stability Cell Atlas^64^. Shared cell types exhibited a remarkable overlap in HRGs. For example, the top ten HRGs of mature B cells in the lung and spleen datasets have a Jaccard index of 0.5, and the top ten HRGs of lung mature B cells and esophageal CD27^+^ B cells have a Jaccard index value of 0.75 (**Table** S3). The HRGs of mature B cells in both the lung and the spleen datasets included canonical markers such as *IGKC, CD79A*, and *MS4A1*, and this consistency extended to CD27^+^ B cells in the esophagus dataset. These results highlight ProtoCloud’s ability to uncover key features of individual cell types, demonstrating the robustness of its explainability across diverse tissue contexts.

### Similarity-guided annotation correction with justification

Reliability is a key aspect of model interpretability, reflecting how trustworthy cell type annotations are. ProtoCloud assesses prediction certainty through multiple complementary approaches. For each cell, our model calculates its similarity score to the closest prototype of the assigned class, enabling quality control of the annotation. These similarity scores between cells and their prototypes exhibit a strong positive correlation with prediction accuracy (**Figure 4**A and **Methods**). We evaluated similarity scores using two calibration metrics (**Methods**), yielding a Brier score^73^ of 0.092 and an Expected Calibration Error (ECE)^74^ of 0.198, indicating that the similarity scores themselves are informative measures of prediction confidence.

**Figure 4:**
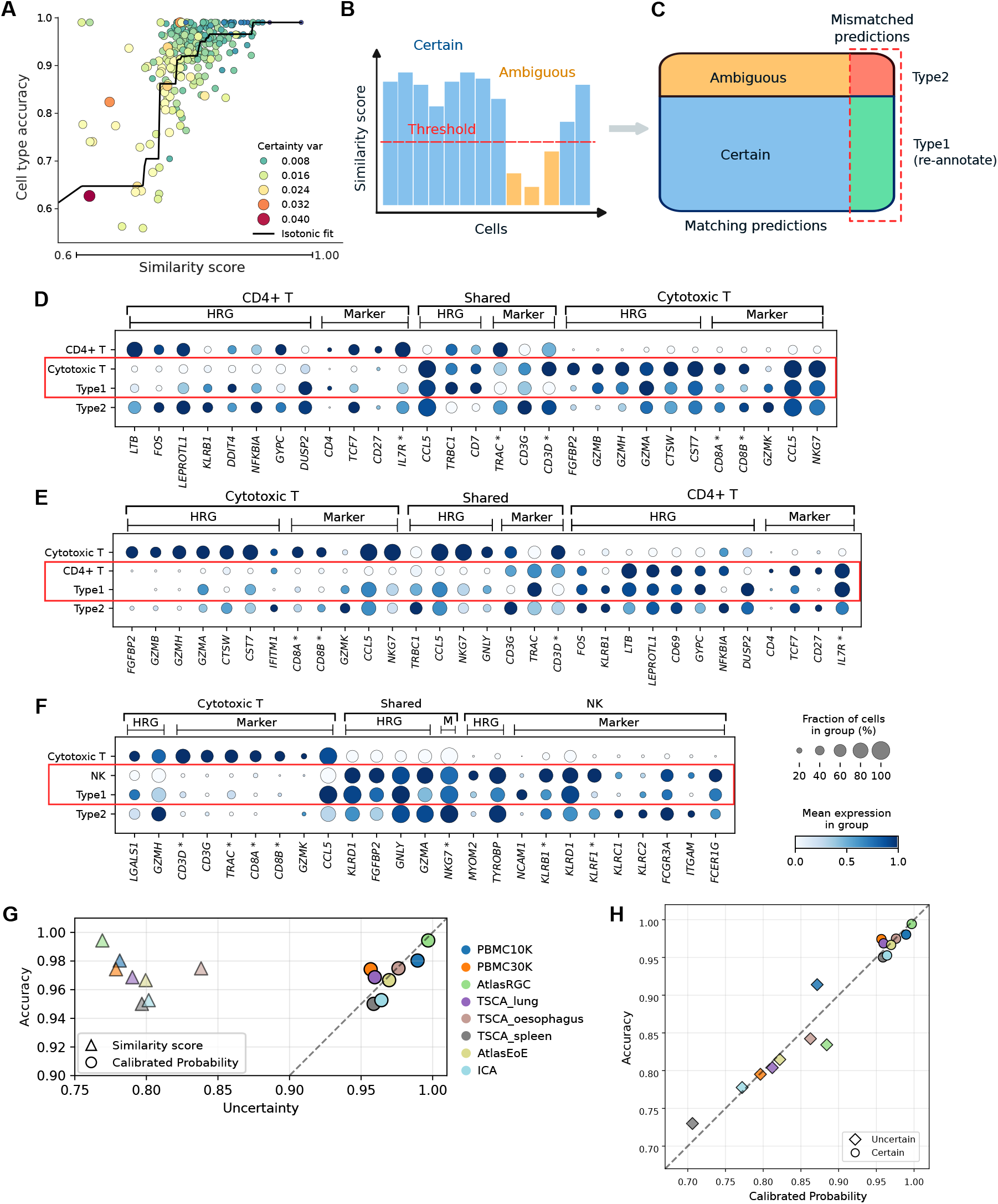
Certainty-based annotation refinement in ProtoCloud. **(A)** The relationship between per-cell type accuracy and similarity scores across eight experimental datasets. The x-axis denotes the similarity score of each cell type, and the y-axis denotes its prediction accuracy. Each point represents a distinct cell type from one of the experimental datasets. Point size and color intensity jointly encode the variance of similarity scores for each cell type. The black curve represents an isotonic regression fit, demonstrating the positive correlation between similarity score and accuracy. **(B-C)** Workflow for annotation refinement in ProtoCloud. **(B)** Step 1: Assign prediction certainty by classifying each cell prediction as “certain” (blue) or “ambiguous” (orange) using class-specific thresholds derived from the training data. Cells with a similarity score above the threshold are classified as “certain”, while those below the threshold are categorized as “ambiguous”. **(C)** Step 2: If the original annotations are available, we can re-annotate confidently predicted cells by comparing predictions with the original annotations. Type 1 cells are cells with high prediction certainty that do not match the original labels (green) and are therefore re-annotated. Type 2 cells (red) are ambiguous cells and have unaligned labels that will not be re-annotated. **(D)-(F)** Expression patterns of major Type 1 annotation pairs in PBMC30K, comparing original annotations (first row), predicted types (second row), Type 1, and Type 2 cells. Type 1 cells should align more closely with the predicted cell type than with the original annotation. Type 2 cells remain ambiguous, showing limited separation between the original and predicted labels. The shared region contains genes that are top-ranked HRGs or known markers in both cell types. Asterisks denote marker genes that are also present in the respective HRG set. **(D)** CD4^+^ T cells versus Cytotoxic T cells. Type 1 cells were originally labeled as CD4^+^ T cells and reannotated as Cytotoxic T cells. **(E)** Cytotoxic T cells versus CD4^+^ T cells. **(F)** Cytotoxic T cells versus natural killer cells. HRG: highly relevant genes, M: Marker. **(G)** Reliability diagram of similarity scores and calibrated similarities of benchmarking datasets. The x-axis of each triangle represents the average similarity score of a dataset, while that of a circle represents the average calibrated probability. The diagonal dashed line indicates perfect calibration, where predicted uncertainty precisely matches observed accuracy rates. **(H)** Reliability diagram of calibrated probability between dichotomous certainty groups of benchmarking datasets. Each point represents one dataset, stratified into “certain” (circles) and “ambiguous” (diamonds) subsets. The dashed diagonal indicates perfect calibration, where predicted certainty matches the empirical accuracy.

From these observations, ProtoCloud categorizes its predictions based on the similarity score into two classes: certain and ambiguous (**Figure 4**B). The similarity threshold used for this classification is tailored to each individual cell type (**Methods**). Predictions with scores exceeding their respective thresholds are classified as “certain”, indicating high confidence in these predictions. Conversely, predictions falling below the threshold are marked as “ambiguous”, signaling the need for cautious interpretation. Given ground truth annotations, ProtoCloud further identifies two distinct types of annotation discrepancies, characterized by prediction certainty and inconsistencies between the model’s prediction and the original annotation (**Figure 4**C). We illustrate each scenario with explanatory examples from the PBMC30K dataset (**Figures 4**D–**4**F):

#### Type 1: Mis-annotated

ProtoCloud confidently predicts a cell type that differs from the dataset’s original annotation. In these cases, the cells are likely mis-annotated in the original dataset and should be re-annotated according to ProtoCloud’s prediction. The gene expression of HRGs for Type 1 cells aligns closely with the predicted cell type rather than the original annotation (**Figures 4**D–**4**F), providing strong evidence for annotation errors in the dataset. For example, in the PBMC30K dataset, cells originally annotated as CD4^+^ T cells but reclassified by ProtoCloud as cytotoxic T cells exhibited an average *NKG7* expression of 3.248, compared to only 0.932 in the originally annotated CD4^+^ T cells and 3.809 in cytotoxic T cells (**Figure 4**D). This substantial increase in *NKG7* expression supports the reclassification. Furthermore, the average Pearson correlation of gene expression between the re-annotated cells and originally annotated CD4^+^ T cells was 0.695, while the correlation with cytotoxic T cells was 0.831, further supporting ProtoCloud’s reclassification.

#### Type 2: Ambiguous mis-prediction

This category includes ambiguous cells whose predicted labels do not align with the original annotations. The expression patterns of the top HRGs for these cells do not distinctly characterize specific cell types. Although the similarity scores may favor one cell type, our model lacks sufficient evidence from key decision-making genes to support a confident prediction. Given their potential to introduce noise into downstream analyses, these cells are excluded from re-annotation.

To provide granular, per-cell certainty estimates, we map the similarity scores to calibrated probability scores using isotonic regression^75^ (**Methods**). These calibrated scores achieved substantially improved performance on both calibration metrics, with the Brier score reduced to 0.048 (from 0.092) and ECE to 0.031 (from 0.198) (**Figure 4**G). Consistent with our dichotomous certainty categorization, cells designated as “certain” exhibited a markedly higher average calibrated score (0.960) compared to those classified as “ambiguous” (0.779). Importantly, observed accuracy aligned with these confidence estimates (0.960 and 0.783), demonstrating that the calibrated probability scores reliably reflect true classification performance (**Figure 4**H).

### ProtoCloud supports time-course data annotation

To further demonstrate ProtoCloud’s ability to assess prediction uncertainty and leverage this functionality for downstream data analysis, we used ProtoCloud to analyze a challenging time-course dataset of mouse retinal ganglion cells (RGCs) following optic nerve crush (ONC)^54^. The cells were collected at seven time points: 0, 0.5, 1, 2, 4, 7, and 14 days post-crush (dpc). The initial model was trained on the control group (0 dpc, **Figure 5**A). The trained model was then applied to the data from the subsequent time point (0.5 dpc), generating predictions categorized as “certain” or “ambiguous”. Cells with “certain” predictions constituted the new training set, using their predicted labels for subsequent training iterations, while ambiguous cases were temporarily held out for validation and re-annotation by the updated model. The initial ProtoCloud model was subsequently trained for 30 epochs on these certain cells and applied to cells from 1 dpc to obtain “certain” and “ambiguous” cells. This iterative training and annotation process was repeated for each subsequent time point.

**Figure 5:**
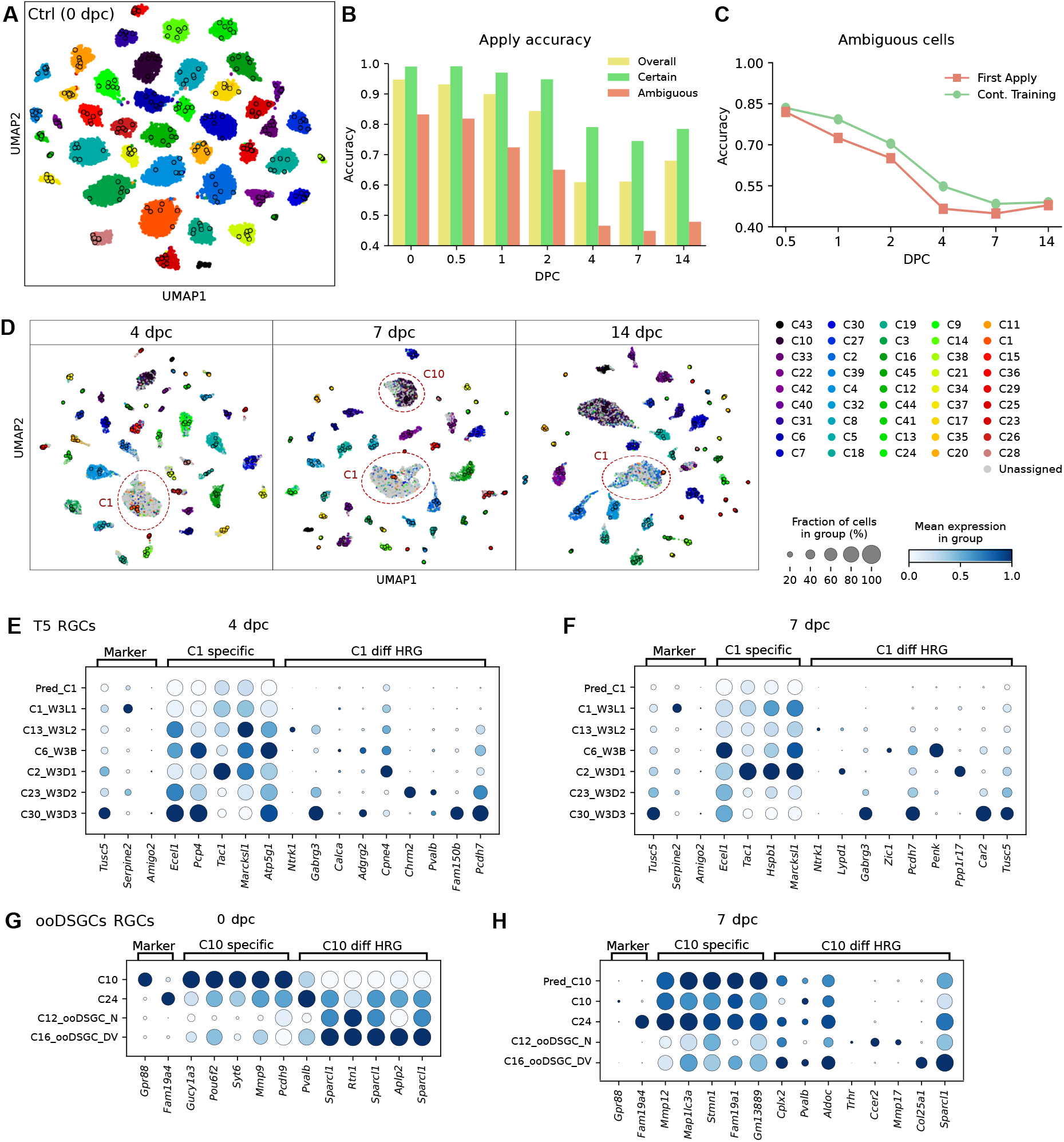
Continue training on the time-course RGC ONC dataset. **(A)** UMAP visualization of the control group (0 dpc) RGCs. **(B)** Initial prediction accuracy at different time points when applying models trained on the previous time point data. For each time point, accuracies are shown for overall predictions (yellow), certain predictions (green), and ambiguous predictions (orange). **(C)** Comparison of prediction accuracy for ambiguous cells only, before and after continued training. “First Apply” shows the accuracy of ambiguous predictions from the initial model application (orange bars in **(B)**). “Cont. Training” shows improved accuracy for these same ambiguous cells after training the model for an additional 30 epochs on cells with certain predictions. **(D)** UMAP visualization of later time points at 4 dpc (left), 7 dpc (middle), and 14 dpc (right). Cells with certain predictions from both the initial application and after continued training are plotted to visualize the results. The prototypes are represented by dots with black edges. Cell types are ordered by injury survival, from the most resilient (C43) to the most susceptible (C28), with an increasing proportion of unassigned cells after 4 dpc. **(E)** HRG expression of T5-RGCs at 4 dpc, and **(F)** 7 dpc. Comparison of top rank HRGs and differentially ranked HRGs between C1 RGCs and other T5 RGCs. ‘C1 specific’ are highly ranked HRGs within C1 (rank ¡ 30) with low relevance in other types (mean rank ¿ 200). “C1 diff HRG” represents genes with the opposite pattern. Re-assigned “unassigned” cells predicted as C1 are included with ‘Pred C1’. ‘Marker’ refers to known canonical markers for each cell type based on established literature^54^. Genes include known markers (*Tusc5* shared by all T5 RGCs; *Serpine2* and *Amigo2*, C1 RGC marker genes), HRGs shared by all T5 RGCs, and differentially ranked HRGs between C1 and other T5 RGCs. **(G)** HRG expression of ooDSGCs-RGCs at 0 dpc and **(H)** 7 dpc including predicted C10 cells (Pred C10). Shared and distinct HRGs are compared between C10 and other ooDSGCs RGCs, with known markers *Gpr88* (a C10 RGC marker) and *Fam19a4* (a C24 RGC marker).

By capturing temporal dynamics in gene expression patterns, ProtoCloud demonstrated its ability to track changes in cellular states throughout the course of injury progression. For confidently annotated cells, the model achieved over 94% accuracy at time points before 4 dpc and maintained accuracy above 68% even at 14 dpc (**Figure 5**B and **Table** S4). Notably, the iterative training strategy improved ProtoCloud prediction accuracy for the initially ambiguous cells by 2-10% (**Figure 5**C). In contrast, both scANVI and CellTypist’s performance declined substantially when using the same limited set of annotated cells for training (**Table** S4).

The proportion of “unassigned” cells increased over time in the original annotations, particularly after 4 dpc, where it increased sharply to approximately 23%. ProtoCloud confidently reclassified 30% to 70% of these previously unassigned cells (**Figure 5**D and **Table** S5). For example, cluster 1 (C1) RGCs, a T5 RGC subtype characterized by marker gene *Tusc5* ^54^, constituted the largest portion of these re-annotated cells. As a result, the estimated survival rate of this subtype increased from 1.2% to 5.6% based on our re-annotations, compared to the original labels. Multiple lines of evidence support the validity of these classifications. The UMAP visualization shows a strong alignment of newly predicted C1 cells with C1 prototypes (**Figure 5**D), and a comparative analysis according to T5 RGC markers and HRGs confirmed that the predicted and reference C1 cells shared highly similar expression patterns (**Figures 5**E–**5**F, Figures S4A–S4B). These predicted C1 cells can be distinguished from other closely related T5 RGCs by HRGs. For example, for the cells at 4 dpc (**Figure 5**E), C1 and predicted C1 cells did not express *Ntrk1* (mainly expressed by C13 RGCs), *Chrm2* (mainly expressed by C23 RGCs), *Gabrg3, Cpne4*, and *Pcdh7* (expressed highly in other T5 RGCs compared to C1 RGCs). The top C1 HRGs showed a systematic but moderate down-regulation pattern in predicted C1 cells compared to reference C1 cells, consistent with an injury-responsive state rather than a distinct cell type (Figures S4C–S4D).

The next largest portion of re-assigned cells belonged to cluster 10 (C10) RGCs, an ooDSGCs RGC subtype distinctly marked by *Gpr88* expression (**Figures 5**G–**5**H and **Figures** S4E–S4F). Although the marker gene exhibited distinct expression in the control group, this signal diminished at later time points. The identified HRGs effectively track these molecular changes, providing reliable references for cell classification throughout the damage progression. At 7 dpc, C10 and predicted C10 cells did not express *Fam19a4* (expressed by C24 RGCs), *Ccer2* and *Mmp17* (mainly expressed by C12 RGCs), and expressed *Col25a1* and *Sparcl1* at a lower level compared to C16 RGCs (**Figure 5**H). At other time points (i.e., 4 dpc and 14 dpc), C10 and predicted C10 cells were highly similar to each other yet clearly distinct from other ooDSGCs RGCs (Figures S4E–S4F). Again, we showed that the predicted C10 cells were close to C10 cells in the UMAP (**Figure 5**D).

We further validate ProtoCloud’s robust transferability across different techniques by applying the model trained on AtlasRGC to an RGC dataset^76^ generated using the Patch-seq technology. ProtoCloud confidently classified 52% of the cells with an accuracy of 97.6%. We then applied the calibrator trained on AtlasRGC to the Patch-seq RGC predictions. Notably, the calibrated scores remained well aligned with the empirical accuracy on this new dataset (Figures S4G–S4I).

### ProtoCloud transfers labels from the esophageal mucosal cell atlas

We applied ProtoCloud to a recently published esophageal mucosal cell atlas (AtlasEoE^25^), which comprises cells from 37 patient biopsies taken from 22 donors across three conditions: healthy, remission eosinophilic esophagitis (EoE), and active EoE. ProtoCloud achieved an overall validation accuracy of 94.5% and a macro F1 score of 0.877 on this large dataset, which comprises 60 prevalent cell types (n = 393,763) and 12 additional rare cell types exhibiting patient-specific bias^25^ (n = 1,215). In particular, the model achieved 97% accuracy specifically for the 12 rare cell states (**Figure** S5A). In contrast, scANVI and CellTypist exhibited frequent misclassifications, particularly confusing related epithelial populations (Figures S5B–S5C).

As expected, ProtoCloud identified cell type-specific HRGs, with known cell type marker genes ranked highly in terms of their prototypical relevance scores (**Figure** S5D). Notably, for *ALOX15* ^+^ macrophages, *PRDM16* ^+^ dendritic cells (cDC2Cs), and group 2 innate lymphoid cells (ILC2s) — three rare cell types of potential importance for EoE pathogenesis—their top HRGs included their respective known marker genes (**Figures 6**A–**6**C). ProtoCloud accurately identified *MMP12* and *ALOX15* as key relevant genes for *ALOX15* ^+^ macrophages (with relevance scores of 0.615 and 0.267, respectively) but not for other macrophages (relevance score *<* 0.03) (**Figure 6**A). *PRDM16* ^+^ cDC2Cs can be distinguished from cDC2Bs based on *PRDM16, TGM2*, and *PIGR* (**Figure 6**B). ILC2, a rare population enriched in active EoE patients^77^, can be distinguished from other IL-13 and IL-5 expressing immune cells (e.g., T helper 2 cells (T_H_2)) by its upregulation of prostaglandin-related genes *PTGDR2, PTGS2*, and *HPGD* ^25^ (**Figure 6**C and **Table** S6). We also identified shared HRGs associated with cell cycle activity (*TUBA1B, HMGB2, TUBB, H2AFZ*, and *KIAA0101*) across proliferating cell populations (**Figure 6**D).

**Figure 6:**
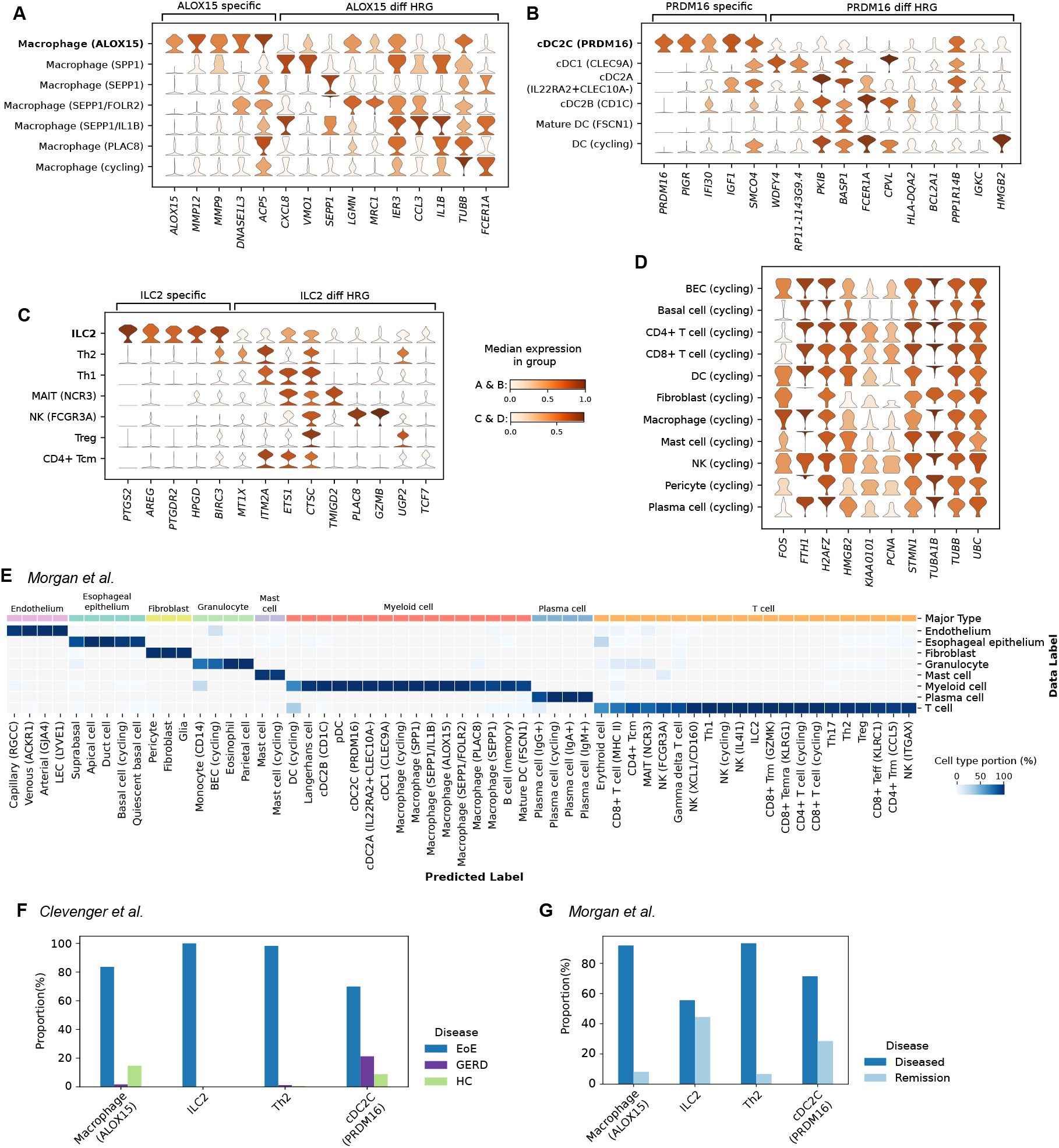
Annotation transfer across EoE datasets and disease conditions. **(A)** Expression profiles of differentially ranked HRGs that distinguish *ALOX15* ^+^ from *SPP1* ^+^ macrophages. ‘ALOX15 specific’ denotes highly ranked HRGs in *ALOX15* ^+^ macrophages (rank ¡ 30) and low relevance in others (mean rank ¿ 200). “*ALOX15* diff HRG” represents HRGs that are highly ranked in contrasting cell types, but not *ALOX15* ^+^ macrophages. **(B)** Expression profiles of differentially ranked HRGs between *PRDM16* ^+^ dendritic cells and *CD1C* ^+^ cDC2B cells. **(C)** Expression patterns of top ILC2-specific HRGs across immune cell populations. Genes were compared between ILC2s and T_H_2 cells. **(D)** Expression patterns of cycling-associated HRGs across proliferating cell populations. **(E)** Annotation transfer correspondence between the AtlasEoE^25^ dataset (72 cell types) and the Morgan et al. dataset^78^ (eight cell types). Color bar: column-wise proportion. **(F)** Cell type composition across disease conditions in the Clevenger et al. dataset^47^ and **(G)** the Morgan et al. dataset. EoE: eosinophilic esophagitis, GERD: gastroesophageal reflux disease, HC: healthy.

We transferred this granular annotation (72 cell states) to two independent EoE datasets^47,78^ (**Figure** S6A), produced by the 10x Chromium^3^ and the SeqWell^6^ platforms, respectively. In the Clevenger et al. dataset^47^, ProtoCloud classified 88.7% of cells as epithelial populations (including quiescent basal, cycling basal, suprabasal, and apical cells), closely aligning with the published proportion of 86.83%. The eight major cell populations identified by Morgan et al.^78^ provided broader categories under which our more granular subcategory annotations were mapped (**Figure 6**E).

Importantly, the three rare but potentially important cell types in the esophagus of EoE patients were clearly identified in the two test datasets (**Figures** S6B–S6E), despite their absence in the original annotations^47,78^. As expected, *ALOX15* ^+^ macrophages expressed the marker genes *ALOX15* and *MMP12*; *PRDM16* ^+^ dendritic cells expressed *PRDM16, PIGR*, and *TGM2*; and ILC2s expressed *IL13, PTGS2*, and *IL17RB* (Figures S6B–S6C). Moreover, we observed a notable increase in the proportion of these cells annotated by ProtoCloud in active EoE patients compared to healthy or remission individuals (**Figures 6**F–**6**G), consistent with previous findings^25^.

We compared this complex label-transfer scenario to the top two benchmark methods: CellTypist and scANVI. While CellTypist also identified major immune cell subsets, it underrepresented rare populations, such as ILC2s and cycling DCs (**Table** S7). Although scANVI also identified these rare cell states (**Table** S7), and in some cases assigned more cells to these rare cell states, the cells specifically identified by scANVI had lower marker gene signature scores compared to those identified by ProtoCloud (Figures S6D–S6E), indicating low-specificity predictions.

## 3 Discussion

We present ProtoCloud, a self-explaining model that augments the variational autoencoder architecture with cell type-specific prototypes to annotate and analyze scRNA-seq data with enhanced transparency. While a trade-off between performance and explainability is often observed in machine learning^79,80^, ProtoCloud demonstrates that these objectives can be achieved simultaneously, matching or exceeding the performance of state-of-the-art methods. To achieve both explainability and high accuracy, ProtoCloud incorporates a carefully designed architecture, including a decomposed latent space to disentangle cell type information from cell type-irrelevant nuisance factors, specialized loss functions (e.g., the atomic loss), and a two-stage training strategy tailored to this architecture. ProtoCloud’s latent space effectively captures both inter-cell type and intra-cell type variations while disentangling additional biological or technological variations without requiring prior knowledge of these cell type-irrelevant factors.

While current deep learning methods achieve impressive accuracy in cell type annotation, ProtoCloud addresses their intrinsic lack of transparency by incorporating cell type-specific prototypes and prototypical relevance propagation (PRP). To the best of our knowledge, ProtoCloud is the first approach to achieve both cell-level and feature-level interpretability in this context. While the granularity of PRP^57^ is often considered a limitation in computer vision applications^58^, this fine-grained gene-level resolution is particularly advantageous for scRNA-seq analysis. The input (gene)-level resolution of PRP aligns with the functions of individual genes and directly reveals genes critical for model decisions. By identifying the genes that strongly activate specific prototypes, ProtoCloud implements a genomic version of the “this looks like that” decision process^81^.

ProtoCloud achieves explainability in terms of three essential criteria: explicitness, faithfulness, and stability^43,82^. The model demonstrates explicitness through its ability to capture highly relevant genes (HRGs) that correspond to both established and novel cell type markers, enabling the distinction between closely related cell types. Faithfulness is evidenced by the prediction certainty assigned to each prediction, which enables the identification of annotation discrepancies, further supported by the HRG expression patterns in disputed cases. Finally, the stability of the learned prototypes is validated across different sequencing techniques, organs, and disease states. Independent analyses identified highly overlapping sets of HRGs for similar cell types in diverse datasets, spanning different technologies and tissues. These capabilities facilitate practical applications in complex biological contexts, helping uncover key molecular features of cell identities.

The practical utility of ProtoCloud was validated through two challenging disease applications. In a time-course analysis of RGCs, ProtoCloud successfully tracked cellular transitions despite limited reference data. In the EoE analysis, it accurately identified and tracked rare disease-specific cell populations across three datasets with varying disease conditions. In both cases, ProtoCloud’s explainability provided molecular-level evidence for disease-specific cellular dynamics, demonstrating its utility in complex, real-world scenarios.

In summary, ProtoCloud transforms single-cell annotation from a black-box prediction process into an evidence-based analytical framework, providing researchers with evidence-based, actionable insights into cellular identity. The model’s ability to identify cell type-specific genes helps bridge the gap between computational predictions and biological understanding. While ProtoCloud effectively identifies genes crucial for cell type classification, it does not directly elucidate the underlying biological mechanisms or environmental interactions. The causal relationships between HRGs and cell types warrant further systematic investigation. By advancing explainable approaches to cell type annotation, ProtoCloud represents a significant step toward more comprehensive and transparent single-cell analysis.

### Limitations of the study

Currently, ProtoCloud requires an annotated reference dataset for training. This means that in the absence of such data, ProtoCloud cannot be directly applied; however, as technology advances, the availability of high-quality reference datasets continues to increase^11^. Second, in this study, we focused on scRNA-seq data analysis. Extending ProtoCloud to other modalities, such as single cell assay for transposase-accessible chromatin using sequencing (scATAC-seq) data^83^, should be straightforward. Applications of ProtoCloud to multimodal data for learning gene regulatory relationships have not yet been explored and represent a promising direction for future research. Finally, although ProtoCloud can identify novel cell types not present in the training data through its built-in uncertainty estimation, how to incrementally update a pretrained ProtoCloud model to incorporate new cell types remains nontrivial and warrants further investigation.

## 4 Methods

### The ProtoCloud Model

We present the implementation details of ProtoCloud, a self-explainable generative model for cell type annotation. ProtoCloud embeds high-dimensional gene expression profiles of individual cells into a low-dimensional latent space and subsequently classifies cells based on their positions in that space. The model comprises four main components: a probabilistic encoder ***g*** : ℝ^*G*^ → ℝ^*d*^, a probabilistic decoder ***f*** : ℝ^*d*^ → ℝ^*G*^, a prototype matrix **P** ∈ ℝ^(*K*·*M*)*×d*^ and a single-layer linear classifier without the bias term ***h*** : ℝ^(*K*·*M*)^ → ℝ^*K*^ . Here, *G* denotes the number of genes, *d* the dimensionality of the latent space, *K* the number of cell types, and *M* the number of prototypes per type (*M* ≥ 1). The prototype matrix **P** contains *K* · *M* prototypes. For each cell type *k*, **P**_*k*_ ∈ ℝ^*M×d*^ represents its set of *M* prototypes, with its *j*th prototype vector denoted by ***p***_*k,j*_, where *j* indexes the prototypes within cell type *k*. The prototypes reside in the same latent space as the cell embeddings, and are randomly initialized and optimized during training to serve as learnable representations of cell types.

ProtoCloud takes raw Unique Molecular Identifier (UMI) counts of cells and cell type annotations as input, in the form of 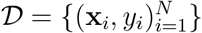 consisting of *N* cell-label pairs, during model training. Each cell *i* is represented by a vector **x**_*i*_ ∈ ℝ^*G*^ of *G* genes assigned to cell type *y*_*i*_ ∈ {1, …, *K*}. We use **y**_*i*_ ∈ {0, 1}^*K*^ to denote the one-hot encoding of the cell type label *y*_*i*_. The probabilistic encoder ***g*** receives log-transformed **x**_*i*_ (specifically, log(1 + **x**_*i*_)) as input, and outputs the parameters ***µ***_***i***_ ∈ ℝ^*d*^ and log(***σ***_***i***_) ∈ ℝ^*d*^ for a normal distribution. The low-dimensional representation **z**_*i*_ for cell *i* is then sampled from this distribution:

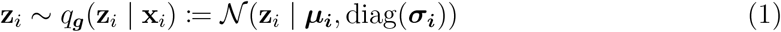

Each low-dimensional embedding **z**_**i**_ serves two purposes: it is used for cell type classification by comparing it with the prototype matrix **P** and feed into classifier ***h***, and it is also passed to the probabilistic decoder ***f*** to reconstruct the UMI counts 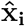.

Single-cell data are noisy, containing both cell type information and frequently capturing cell type-irrelevant nuisance factors such as experimental batches. We thus partition the latent embedding **z**_*i*_ into two parts: the first half, 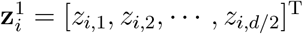, is designed to specifically capture cell type information, while the second half, 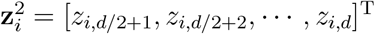, is designed to capture cell type-irrelevant factors. Consequently, only 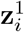 is used for cell type classification, while both 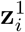 and 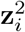 are used by the decoder ***f*** to reconstruct the gene expression profile 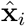.

The classifier ***h*** takes a vector of similarities between cell *i* and the *K*·*M* prototypes in **P**, and produces a probability distribution over the *K* cell types. We compute the similarity between cell *i* and the *j*-th prototype of cell type *k* using a scaled Cauchy distribution density function:

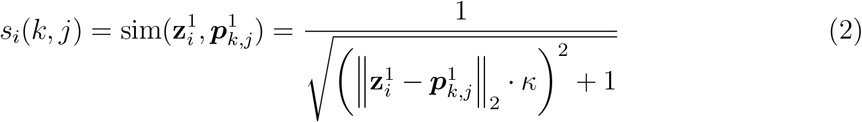

where *κ* is a learnable, positive scalar parameter and 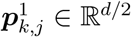 is the first half of ***p***_*k,j*_. This formulation assigns a maximum similarity score *s*_*i*_(*k, j*) of 1 when the embedding 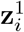 exactly matches prototype 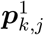, and the similarity smoothly decreases as the Euclidean distance increases. The similarity *s*_*i*_(*k, j*) indicates the influence of prototype *j* of cell type *k* on the classifier prediction, and the maximum similarity can serve as a prediction certainty measurement (see section “Anomaly detection certainty threshold”). The classifier ***h*** takes the vector of similarity scores

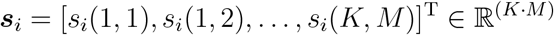

as input and is trained via cross-entropy (CE) loss:

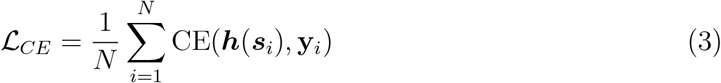

The probabilistic decoder ***f*** is essential for providing cell-level interpretations by reconstructing the gene expression profiles from the latent embeddings. For scRNA-seq data, the UMI count distribution for a gene *g* in cell *i* is generally assumed to follow a negative binomial (NB) distribution^84^. Accordingly, the decoder outputs the mean *m*_*i,g*_ ≥ 0 and the dispersion *r*_*k,g*_ *>* 0 parameters. Given the training data *D* with cell type labels, we thus use a cell type-specific dispersion parameter *r*_*k,g*_ for gene *g*, to capture cell type-specific variability in gene expression. The mean parameter *m*_*i,g*_ for gene *g* remains cell-specific. Specifically, the probabilistic decoder outputs a *G*-dimensional probability vector (non-negative and summing to 1), which is multiplied by the total UMI count *n*_*i*_ for cell *i* to obtain the mean vector ***m***_*i*_. The cell type-specific dispersion *r*_*k,g*_ is learned for each gene *g* and cell type *k*. Thus, the ProtoCloud model likelihood can be written as

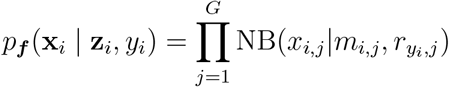

ProtoCloud has a variational autoencoder (VAE) architecture backbone, and we need to calculate the evidence lower bound (ELBO) to optimize the probabilistic encoder (***g***) and decoder (***f***). The ELBO includes a Kullback–Leibler (KL) divergence term between the variational distribution *q*_***g***_(**z**_*i*_ | **x**_*i*_) (Equation 1) and a latent prior *p*(**z**_*i*_). Conventionally, *p*(**z**_*i*_) is a standard multivariate normal distribution. In our case, it becomes complicated because we want cells to be close to their cell-type prototypes. Moreover, we decompose the latent embedding into two components: 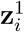 for cell type-specific information and 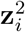 for cell type-independent factors. To achieve these goals, we consider a mixture of VAEs with shared encoder and decoder networks. Each VAE mixture component has a prior centered on one of the *M* prototypes for each cell type *k*. Only the training cells with label *k* are used to calculate the KL-divergences for these *M* VAEs. The KL divergence for each prototype is weighted by its relative similarity within that class, 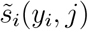^59^.

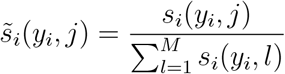

The overall KL divergence is then calculated as the weighted sum of the KL divergences for the cell type-specific and cell type-irrelevant components:

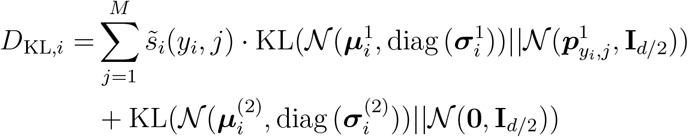

where KL(*q*||*p*) denotes the KL divergence between probability distributions *q* and *p*, 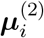 and 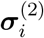 are the second halves of the mean and standard deviation parameters of the latent normal distribution for **z**_*i*_, and **I**_*d/*2_ is the identity matrix with a dimensionality of *d/*2.

Combining these elements, we formulate the VAE objective function to maximize the ELBO:

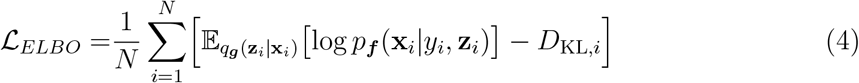

As both the encoder (***g***) and the decoder (***f***) networks are shared across all cell types and prototypes, ProtoCloud remains computationally efficient, comparable to conventional VAEs.

#### 4.0.1 The orthogonal loss to help learn diverse prototypes for each class

Training ProtoCloud using the sum of the cross-entropy loss (Equation 3) and the negative ELBO (Equation 4) frequently encounters the prototype collapse problem, in which the *M* prototypes of cell type *k* converge to a single point, often near the center of the embeddings of class *k* cells. This collapse prevents the model from learning diverse representative cell prototypes that capture the intra-class diversity. To help mitigate this problem, we introduce an extra orthogonal loss

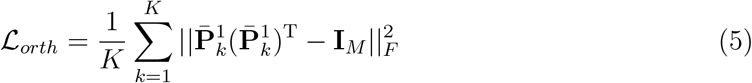

where ∥·∥_*F*_ denotes the Frobenius norm and 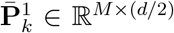 is the first half of the centered prototype matrix for class *k*. Specifically, we compute the mean of the *M* prototypes for class *k*

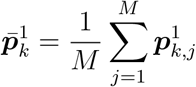

and subtract this mean from each prototype 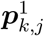 within that class to obtain a centered matrix. By encouraging the rows of 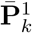 to be orthogonal, this loss helps maintain multiple, distinct prototype directions within each class.

#### 4.0.2 The atomic loss to help learn diverse and accurate prototypes

To further encourage cell embedding to be close to cell type-specific prototypes, we introduce an attractive force. Let mask_*i*_ be a binary vector of length *K*·*M* where mask_*i*_[*M*· (*k* − 1) + *j*] = 1 if the prototype *j* of cell type *k* belongs to the same cell type as cell *i*, and 0 otherwise. The attractive force is defined as:

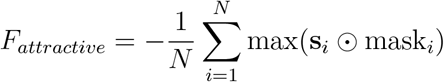

where ⊙ denotes element-wise multiplication. This force encourages each cell embedding 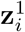 to be close to at least one of its *M* prototypes in its own class. However, a trivial solution could minimize this force by collapsing all cells to the same point, overlapping with all *K*·*M* prototypes.

To counteract such collapse and address potential issues arising from multiple prototypes per class and rare cell types, we add a repulsive force. As ProtoCloud uses multiple prototypes for each class, it’s possible that some prototypes of cell type *k* are close to the embeddings of cells from other classes. Furthermore, in the presence of rare cell types, both their prototypes and the encoder may not be well trained to embed these cells effectively^32^, leading to misplaced prototypes. Such misplacements make it challenging to draw reliable cell-level interpretations. The repulsive force pushes away prototypes of other classes from the cell embeddings of class *k* cells

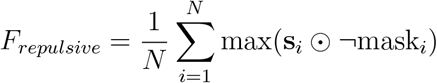

where ¬mask_*i*_ is the negation of the binary mask vector mask_*i*_. Together, the dynamics of attractive force and repulsive force—analogous to inter-molecular forces between atoms—are combined into the atomic loss

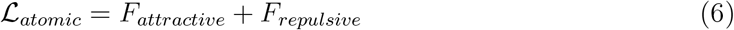

#### 4.0.3 Shaping latent spaces through a two-stage curriculum and latent decomposition

Latent embeddings learned by variational autoencoders can also fluctuate, similar to the latent embeddings learned by conventional autoencoders, as observed by pioneers in deep learning such as Geoffrey Hinton, resulting in latent space fragmentation, where cell embeddings of the same type are widely separated and partitioned into disjoint regions by embeddings of other cell types. This fragmentation is primarily driven by technical factors such as batch effects, donor-specific variation, and sequencing depth, which can artificially separate otherwise biologically homogeneous cell populations. To learn more compact embeddings in which cells of the same cell type form a unified and cohesive region, we employ a two-stage training strategy, along with latent space decomposition, in ProtoCloud.

In the first stage, we train ProtoCloud by minimizing the loss function

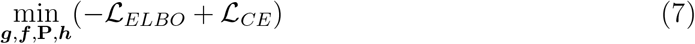

During this stage of training, the prototypes of the same class are encouraged to collapse toward a single vector, thereby drawing the corresponding cell embeddings together to form a unified, compact region around their prototype. Notably, as classification utilizes only the first half of the latent dimensions, batch-related information is encouraged to be captured within the second half of the dimensions, effectively disentangling batch effects from cell type information.

In the second stage of training, we reintroduce prototype diversity through the orthogonality and atomic loss terms. The orthogonal and atomic losses are designed to generate repulsive forces among embeddings, promoting separation in the latent space and increasing the diversity of learned prototypes. The updated loss function in this stage then becomes

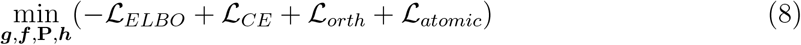

To facilitate feature (gene)-level interpretation of cell type annotation in the second stage, we add a penalty term *L*_1_ to the weights of the first layer of the encoder. This penalty encourages sparsity by driving the weights of unimportant genes for cell type annotation towards zero. The strength of the *L*_1_ regularization is set to the inverse of the number of elements in the first layer weight matrix.

By decoupling the training process into two stages, ProtoCloud progressively refines its representations: the first stage prioritizes inter-class diversity, while the second stage enhances intra-class diversity. After a ProtoCloud model has been trained, it can be used to annotate cells, taking only raw UMI count data of cells as input.

### Prototypical Relevance Propagation for gene-level interpretation

To obtain gene (feature)-level interpretation and identify which genes drive ProtoCloud’s predictions, we use prototypical relevance propagation (PRP)^58^. For each cell type *k* and one of its associated prototypes ***p***_*k,j*_, we first calculate the cell-prototype similarity for a given cell *n* assigned to cell type *k*:

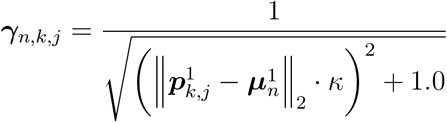

where 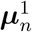 is the first half of the mean vector of the variational normal latent distribution *q*_***g***_(**z**_*n*_ | **x**_*n*_). The similarity vector ***γ***_*n*_ ∈ ℝ^(*K*·*M*)^ is backpropagated through the encoder following the LRP rules. The initial relevance ***R***^(*L*+1)^ is a (*K*·*M*)-dimensional vector, with each element set to −1, except for the *M* elements corresponding to the prototypes of cell type *k*, which are set to either 1 (the prototype closest to cell *n*) or 1*/M* (the other prototypes). The number of hidden layers in the encoder network is *L*.

More specifically, let *o*_*j*_ represent the *j*th output of a fully connected layer *l*, before the layer’s rectified linear unit (ReLU) activation, then

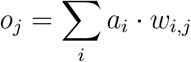

where *w*_*i,j*_ is the weight connecting the *i*th input neuron with activation *a*_*i*_ at layer *l*, to the output neuron *j* at layer *l* + 1. Thus, the relevance from the layer’s output 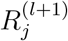 is propagated to its input 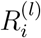, according to the following equation:

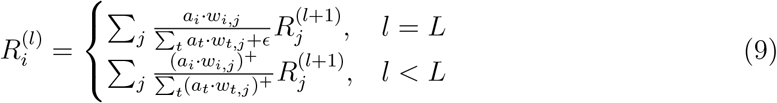

The parameter *ϵ* = 1*e*^−6^ is used by default. For convenience, we use the notation (·)^+^ = max(0,·). We only consider positive contributions to the relevance for the encoder layers because the input gene expression values are non-negative, and we want to find genes that positively contribute to the prediction of cell types. The rules in Equation 9 are called the *ϵ*-rule (when *l* = *L*) and *z*^+^-rule (when *l < L*), or equivalently the *αβ*-rule with *α* = 1 and *β* = 0^57^.

The final relevance scores 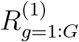 for all the *G* genes are normalized to the range [0, 1] to facilitate direct comparison of relevance scores across different prototypes. To obtain a gene-level relevance score for cell type *k*, we weighted the relevance scores across all *M* prototypes associated with cell type *k* and all training cells with label *k*.

### Data augmentation for training robust ProtoCloud models

Single-cell RNA-Seq data often exhibit class imbalance, with some cell types being extremely rare (e.g., accounting for *<* 0.005% of the recovered cells) and dominant cell types^25^. This disparity is prevalent in data from various biological contexts, such as development, homeostasis, and cancer, posing significant challenges for machine learning algorithms^85^. A common approach to address this imbalance is oversampling, where rare class populations are duplicated to create more balanced training data. However, this simple approach can lead to overfitting of the rare populations, as it does not capture the underlying data distribution of a rare class.

To mitigate this class imbalance problem, we generate artificial cells for rare populations using statistical modeling of the gene count distribution for each gene in each rare cell type. Specifically, for each rare cell type (defined as having a proportion less than 0.5*/K*, where *K* is the number of cell types), we generate artificial samples to ensure that the total number of cells from each rare population reaches 0.5 ×*N/K*, where *N* is the total number of cells in the original dataset. We sample an artificial cell’s UMI count vector 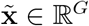 from a multinomial distribution. The distribution’s rate probability vector parameter ***r*** is estimated from a randomly drawn UMI count vector **x** from a cell of the rare population:

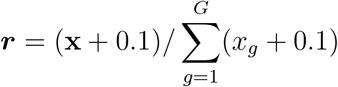

Adding 0.1 to each element of **x** ensures that ***r*** is a dense probability vector that sums to 1. By sampling from this multinomial distribution with probability vector ***r***, we can generate non-zero counts for genes that had zero counts in the original sample **x**. The size parameter (number of trials) for the multinomial distribution is sampled uniformly from the interval:

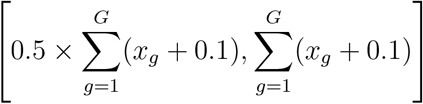

This data augmentation step ensures a more balanced representation of all cell types in the training data, thereby reducing the risk of overfitting rare cell types.

### Model architecture and training

By default, ProtoCloud employs an encoder with three hidden layers (1,024, 512, and 256 neurons, respectively) and a decoder with two layers (512 and 1,024 neurons, respectively). Each layer consists of a bias-free linear transformation followed by batch normalization^86^ and ReLU activation^87^. The model uses a 20-dimensional latent space and maintains six prototypes per class. Training is performed for 100 epochs using the AdamW optimizer^88^ with a learning rate of 1 *×* 10^−3^. The mini-batch size is 128 for datasets with fewer than 100,000 cells and 1,024 for larger datasets. These hyperparameters remain constant across experiments unless otherwise specified.

To estimate the time required to train ProtoCloud for 100 epochs, we down-sampled one dataset to various numbers of cells (10,000, 50,000, 100,000, 200,000, and 300,000 cells). All experiments were conducted on a server with an NVIDIA Tesla V100 16GB GPU on a single node. Training ProtoCloud is efficient, even for relatively large datasets with 300,000 cells when using adaptive batch sizes (128 for smaller datasets with fewer than 100,000 cells and 1,024 for datasets exceeding 100,000 cells). The training times across the different dataset sizes are shown in **Figure** S1A. For large datasets, the training time can be further decreased by reducing the training epochs (default: 100 epochs).

### Ablation study

To systematically assess the contribution of each component in ProtoCloud, we conducted an ablation study examining three key design choices: the disentangled latent space, the two-stage curriculum, and the inclusion of two auxiliary losses (orthogonal and atomic losses). These studies evaluate model performance through multiple complementary perspectives, including batch effect separation, prototype embedding distributions, and similarity score ranges.

We evaluate the quality of prototype embeddings using Euclidean distances in the first half of the embedding space. Our evaluation differs from conventional clustering metrics such as the silhouette coefficient, which prefer large inter-class distances and small intra-class distances. Instead, we require both distances to be substantial to effectively capture cell-type distinctions while preserving within-type biological variation.

#### Disentangled vs. unified latent space

Our standard architecture models the first half of the latent dimensions of a cell with a multivariate normal distribution that is constrained to be close to its prototypes and uses these dimensions for similarity score computation, while the remaining dimensions are encouraged to follow a standard multivariate normal distribution. The unified variant removes this architectural separation in the KL divergence, encouraging all latent dimensions of a cell to follow a multivariate normal distribution that is constrained to be close to its prototypes and using all latent dimensions for the similarity calculation.

#### Two-stage vs. single-stage curriculum

In the standard two-stage curriculum, ProtoCloud is initialized and trained with only the baseline VAE losses, ℒ_*ELBO*_ and ℒ_*CE*_, for the first 30 epochs. The model then enters the second stage after 30 epochs, where the atomic loss and orthogonal loss are added. The single-stage training baseline incorporates all loss components from initialization, training end-to-end for 100 epochs.

#### Loss function component ablation

We evaluated three loss configurations against our standard formulation. The baseline VAE configuration uses only ℒ_*ELBO*_ and ℒ_*CE*_; the atomic-only version adds the atomic loss, ℒ_*atomic*_, to the baseline VAE losses; and the orthogonal-only version adds the atomic loss, ℒ_*orth*_, instead.

### Reliability and uncertainty estimation

ProtoCloud uses similarity scores as a more interpretable and robust measure of uncertainty than softmax probabilities, which often saturate toward extreme values^89^. We quantify prediction confidence using the maximum similarity score between each cell’s first-half latent embedding and the prototypes.

Dichotomous certainty thresholds are determined at the end of training based on the training data. These thresholds are computed separately for each cell type, set at the 10th percentile of similarity scores between training cells and their corresponding class prototypes. Predictions with similarity scores exceeding their respective cell type thresholds are marked as “certain”, indicating strong confidence in the assigned annotation. Certain predictions that disagree with the original labels are categorized as Type 1 classifications. Conversely, predictions below this threshold are marked as “ambiguous”. Ambiguous predictions that disagree with the original labels are categorized as Type 2 classifications. As we adopt a conservative threshold, “ambiguous” reflects insufficient confidence for re-annotation rather than incorrect predictions.

To further improve reliability and interpretability, we transformed similarity scores into calibrated certainty scores through an adaptive calibration approach. The test data were randomly partitioned into two subsets: 25% were used to fit a calibrator using ProtoCloud predictions, similarity scores, and corresponding cell type labels, while the remaining 75% were used to validate the resulting calibrated scores. First, we trained a global isotonic regression^75^ calibrator on the entire dataset to establish a baseline certainty mapping, which outputs a calibrated probability for each prediction. For rare cell types (see Data Augmentation), the global calibration model was employed to prevent overfitting and ensure stable certainty estimates. For cell types with sufficient samples, we further trained cell type-specific calibrators to map similarity scores to calibrated certainty values.

### Differentially ranked highly relevant genes

The genes for each cell type are ranked according to their relevance scores from high to low. Differentially ranked HRGs between two cell types are identified using two criteria: (1) ranked top *t* (default 20) in exactly one of the interested cell types, or (2) showing a ranking difference greater than *u* (default 200) between the two cell types.

### Baselines for comparison

#### Seurat v4^27^

We used its reference-based label transfer functionality, following the workflow described in the Seurat integration tutorial (https://satijalab.org/seurat/articles/integrationmapping). Specifically, we employed the *FindTransferAnchors()* function with the projected PCA option to identify correspondences between reference and query datasets, followed by the *TransferData()* function to propagate cell type labels to the query dataset.

#### scANVI^26^

We utilized the *scvi*.*model*.*scANVI* API from the Python package scvi-tools (v1.1.5). Following best practices, we first trained an scVI model on the dataset for a maximum of 400 epochs. The resulting pre-trained scVI model was then used to initialize the scANVI model, which was subsequently trained for an additional 20 epochs to refine the annotations. For the time-course RGC ONC data analysis, the initial model was trained on labeled control data and used to predict labels for the cells from the subsequent time point. These predictions were then incorporated into the previous training set to train the next version of the model. All predictions were added to the training set as the model lacks a confidence output. This process was repeated sequentially through all developmental stages.

#### CellTypist^28^

CellTypist models were trained using the SGD algorithm with a maximum of 500 iterations. For the time-course RGC ONC data analysis, we used CellTypist with an iterative training approach: the initial model was trained on labeled control data and used to predict labels for the cells from the subsequent time point. Predictions with corresponding confidence scores greater than 0.5 were then incorporated into the previous training set to train the next version of the model (the fraction of confident predictions was small; **Table** S4). This process was repeated until the cells from every time point had been predicted.

#### TOSICA^60^

We implemented TOSICA using its standard workflow. For all datasets involving human cells, the pre-prepared human GO (‘human gobp’) mask was used. For the AtlasRGC dataset, the corresponding mouse GO mask (‘mouse gobp’) was used.

#### SIMS^62^

Following the authors’ recommendation for typical convergence, the model was trained for 10 epochs on each dataset. All other parameters were maintained at their default values.

#### scPoli^**61**^

The method requires batch information as input data. The model was trained for a total of 100 epochs, which included an initial 40 epochs of pre-training, as specified.

#### scGPT^40^

Due to the significant computational complexity associated with fine-tuning large foundation models, we employed scGPT in a zero-shot inference setting. This approach leverages the pre-trained whole-human scGPT model to generate cell embeddings and annotations for our query datasets without any additional training. For the AtlasRGC dataset, genes were first mapped to human orthologs prior to analysis.

#### scBERT^38^

We utilized the pre-trained scBERT model and fine-tuned it on our labeled reference data for 10 epochs, following the standard pre-train and fine-tune paradigm of the method. Due to scBERT’s computational complexity and gene constraints from its pretrained model, we obtained results only for the PBMC10K, PBMC30K, AtlasRGC, TSCA lung, and TSCA oesophagus datasets.

### Evaluation metrics

We evaluated the performance of cell type classification using three standard metrics: accuracy, macro F1-score, and Cohen’s Kappa coefficient^65^. Accuracy offers an overall view of model performance, macro F1-score is more sensitive to minority classes by treating each class equally, and Cohen’s Kappa coefficient (*κ*) quantifies the agreement between predicted and true labels. More specifically, *κ* corrects for chance agreement:

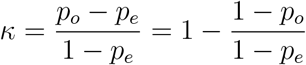

where *p*_*o*_ is the classification accuracy, or the empirical probability of agreement on the label assigned to a cell, and *p*_*e*_ is the expected classification accuracy when both true labels and predicted labels are randomly assigned to cells. Thus, Cohen’s Kappa coefficient is a function of the classification accuracy. The advantage of *κ* over accuracy is that *κ* is sensitive to rare classes. We used scikit-learn (v1.5.1)^90^ to calculate *κ*. Together, these three metrics (with higher values indicating better performance) ensure reliable and fair assessment across all cell types, including rare populations.

To assess batch separation in the latent space, we used the batch entropy score (BES) and the scib-metrics framework (v0.5.1)^91^. BES quantifies batch mixing based on the composition of *k*-nearest neighbors (*k*-NN). Let *B* be the total number of batches, *p*_*i,b*_ represent the proportion of the neighbors of cell *i* belonging to batch *b*. We then averaged the entropy across all cells to obtain the overall metric:

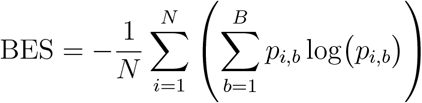

where *N* is the total number of cells. A BES value close to log(*B*) indicates greater batch mixing and weaker batch effects, whereas a lower value (with a minimum of 0) suggests stronger batch separation within the latent space.

Additionally, we applied the scIB metrics to evaluate biological conservation and batch correction in the latent space, focusing on two key dimensions: biological conservation and batch correction. Biological conservation was evaluated using five complementary metrics: Normalized Mutual Information (NMI) and Adjusted Rand Index (ARI) (computed using K-means clustering) to measure cluster agreement; the isolated label F1 score to assess the preservation of rare cell types; and cell-type Adjusted Silhouette Width (ASW) along with cell-type LISI (cLISI) to quantify neighborhood purity. Batch correction performance was assessed using batch ASW, integration LISI (iLISI), and the k-nearest neighbor batch effect test (kBET) to measure local mixing; graph connectivity to evaluate the structural integrity of the neighbor graph; and principal component regression (PCR) comparison to quantify the variance explained by batch covariates.

To assess the biological relevance of the top-ranked HRGs, we employed three complementary metrics: gene signature scoring, cell type specificity quantification, and pathway enrichment analysis. To show whether the HRGs identified by ProtoCloud capture cell type-specific transcriptional patterns, we calculated the gene signature score for each cell type’s top HRGs across all cells. This analysis was performed using the *sc*.*tl*.*score genes* function from the Scanpy Python library^92^. To further investigate the gene-level specificity of HRGs, we compared the top-ranked HRGs against a curated set of known markers from scType^31^. We used the tau index (*τ*), a widely adopted metric for measuring tissue or cell type specificity in transcriptomic studies^72^. The tau index is defined as:

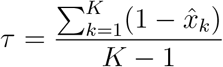

where *K* denotes the number of cell types, and 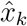 represents the average expression level of a gene in cell type *k*, normalized by the maximum average expression across all cell types. The tau index ranges from 0 to 1, where *τ* = 0 indicates ubiquitous expression across all cell types, and *τ* = 1 indicates highly specific expression that the gene is strongly expressed in only one cell type and is nearly absent from all others. To show the biological functions associated with HRGs, we performed Gene Ontology (GO) enrichment analysis^93^ targeting biological processes (BP). Specifically, we utilized the *GO Biological Process 2025* library^94^ to query the HRGs identified for each cell type. Enrichment significance was assessed using Fisher’s exact test, followed by the Benjamini-Hochberg^95^ procedure to adjust the *p*-values for multiple comparisons (False Discovery Rate, FDR). We reported the top enriched biological processes for each cell type to characterize the functional relevance of identified HRGs.

To evaluate the extent to which the certainty estimates are calibrated, we employed two metrics: the Brier score^73^ and the expected calibration error (ECE)^74^. The Brier score quantifies the overall probabilistic accuracy by calculating the mean squared error between the predicted certainties and the actual outcomes:

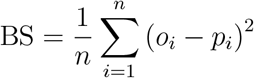

where *o*_*i*_ is the outcome of the prediction at instance *i* (1 if the prediction is correct, otherwise 0), and *p*_*i*_ is the predicted probability. To specifically measure the discrepancy between the predicted confidence and empirical accuracy, we utilized the ECE. This metric partitions the predictions into a number of bins (*B* = 20 by default) based on their confidence scores. The goal is to calculate the weighted average of the absolute difference between the mean predicted confidence and the fraction of positive outcomes within each bin:

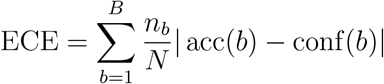

where *B* is the number of bins, *n*_*b*_ is the number of samples in bin *b*, acc(*b*) is the accuracy, and conf(*b*) is the average model confidence (e.g., predicted similarities) within the bin. While both metrics assess calibration, they capture distinct aspects: the Brier score emphasizes the accuracy of probability estimates, whereas the ECE emphasizes the alignment between confidence and correctness. Both metrics range from 0 to 1, where a value of 0 indicates perfect calibration.

### Data collection and processing

For data preprocessing, mitochondrial and ribosomal protein-coding genes are removed from all datasets. When applying the trained ProtoCloud models to unseen data (EoE studies; see below) or continuing training with a new dataset (RGC ONC studies; see below), only the genes shared with the data used for training the pre-trained ProtoCloud models are utilized. Missing genes in the test data are filled with zeros.

#### PBMC10K

We used the preprocessed PBMC dataset^3^ downloaded from scvi-tools^96^. This dataset includes 12,039 cells, 3,346 genes, and two batch groups, spanning nine annotated cell types. No filtering was applied except for removing the cells labeled as “unknown”.

#### PBMC30K

This PBMC dataset was derived from two experiments, each employing multiple scRNA-seq methods^63^. The original dataset consisted of 30,495 cells across ten cell types. We filtered out cells with fewer than 1,000 counts and genes with fewer than 200 total counts. The “unassigned” cells were removed from the data, resulting in 14,978 cells that passed filtering, with the top 3,000 HVGs selected for analysis^92^.

#### AtlasRGC

An atlas of mouse retinal ganglion cells (RGCs) in the adult retina^54^. The dataset comprises three batches and a total of 45 RGC subtypes, with cell proportions ranging from 8.4% (C1) to 0.15% (C45). We excluded cells with fewer than 1,000 total counts and cells labeled as “unknown”, resulting in 35,699 cells. Genes with a total count below 500 were removed, and the top 3,000 HVGs were selected for analysis.

#### Time-course RGC Datasets

The time-course RGC dataset, profiling retinal ganglion cells following optic nerve crush, was obtained from the same study^54^ as AtlasRGC. This dataset includes varying numbers of cells at each time point (**Table** S4), with “unassigned” cells after 0 days post-crush (0 dpc). In our experiment, only cells from 0 dpc were used to train the initial model, based on the top 3,000 HVGs identified from the 0 dpc data. Cells from subsequent time points were then incorporated using the same set of 3,000 HVGs.

#### Patch-seq RGC

The raw single cell RNA-Seq data collected from Patch-seq technology, featuring 472 cells and 55,542 genes^76^. We applied the ProtoCloud model trained on AtlasRGC to this dataset. The top 3,000 HVGs from AtlasRGC were used, of which 2,924 were present in the Patch-seq RGC dataset.

#### TSCA Datasets - Lung, esophagus, and spleen

These datasets are from three human primary tissues, each exhibiting distinct levels of sensitivity to ischemia. The spleen, considered the most stable, contains 94,257 immune cells spanning 30 cell types. The esophageal mucosa includes 87,947 cells from 19 cell types, while the lung, identified as the least stable, comprises 57,020 cells across 28 cell types. For all three datasets, we filtered out cells with fewer than 1,000 counts and removed genes with fewer than 500 total counts. The top 3,000 HVGs were selected for analysis for each dataset. For comparative models requiring batch information, patient identity was used as the batch variable, resulting in five, six, and five batches for the spleen, esophageal mucosa, and lung datasets, respectively.

#### ICA

The cross-tissue Immune Cell Atlas dataset is part of the Human Cell Atlas project^8^. It includes over 300,000 immune cells, isolated from 16 different tissues obtained from 12 deceased adult donors. Organ origins are used when batch information is needed by comparative models. Cell identities were annotated using CellTypist into 43 cell subtypes. The top 3,000 HVGs were selected for analysis.

#### AtlasEoE

The esophageal cell atlas^25^ consists of 421,312 scRNA-seq profiles. These samples were taken from 15 EoE patients (eight in active disease state and seven in remission) and seven healthy participants. The dataset includes 60 prevalent cell types grouped into major categories (e.g., epithelial cells, stromal cells, monocytes/mac/DC cells, B cells, T/NK/ILC cells). Additionally, the atlas has 12 rare cell subsets, comprising 1,215 cells in total. For methods requiring batch information, donor identifiers were treated as batch labels, resulting in 22 batches. For this dataset, we used the highly variable genes (*G* = 2, 956) provided by the authors.

#### Clevenger et al. Dataset

This dataset^47^ comprises 151,519 esophageal cells from six healthy donors, six patients with EoE, and four patients with gastroesophageal reflux disease (GERD). The authors identified eight major cell populations and focused on epithelial cells in their study. We applied the ProtoCloud model, trained on the AtlasEoE data, to this dataset.

#### Morgan et al. Dataset

This dataset^78^ comprises 14,242 esophageal cells collected from pediatric EoE patients in active or remission EoE, encompassing eight major cell types. We applied the ProtoCloud model, trained on the AtlasEoE data, to this dataset.

## Acknowledgments

We thank Dr. Anne Condon, Jeffrey Niu, Minuk Ma, and Carlos Vasquez for their constructive criticism and valuable feedback on this work, and the members of the Ding group for insightful discussions and support. This work was supported by a Discovery grant from the Natural Sciences and Engineering Research Council (NSERC) of Canada, and a department startup fund from the University of British Columbia (to J.D.). J.D. is a Canada Research Chair and is supported by the Canadian Institutes of Health Research through the Canada Research Chair Program. The computational resource is partially supported by the Canada Foundation for Innovation & John. R. Evans Leader Fund (to J.D.). This research was supported in part through the computational resources and services provided by Advanced Research Computing at the University of British Columbia.

## Author’s contributions

J.D. conceived the project. J.D. and K.G. developed the model. K.G. conducted experimental analyses with guidance from J.D. K.G. and J.D. interpreted the results and wrote the manuscript.

## Supplementary Figures

**Figure S1:**
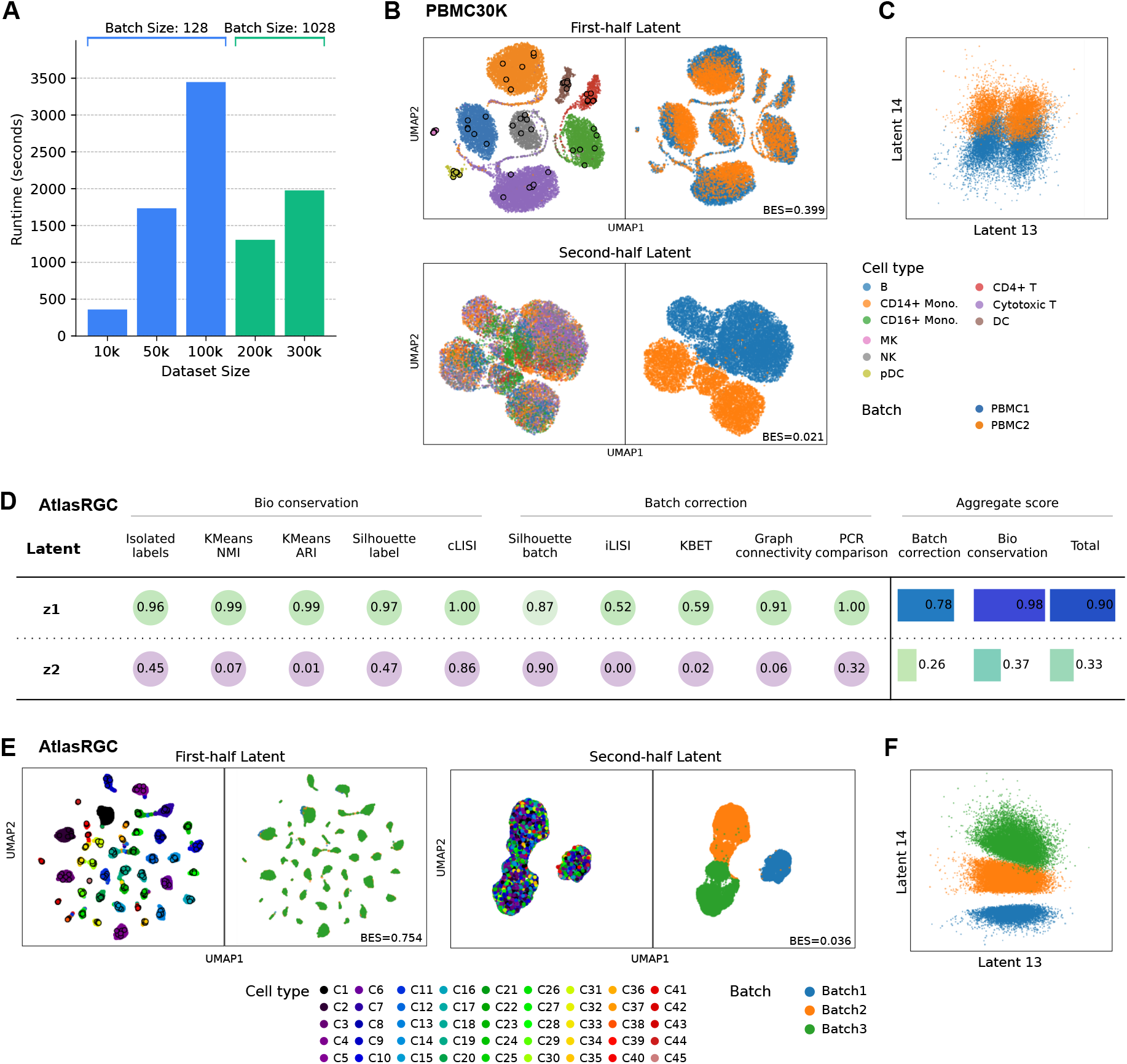
ProtoCloud runtime and visualization of latent embeddings, related to Figure 3. **(A)** Training time of ProtoCloud versus data size. All experiments were conducted on a server with one NVIDIA Tesla V100 16GB GPU on a single node. Training time scales linearly with dataset size, with smaller datasets processing efficiently and larger ones benefiting from an increased batch size. **(B)** UMAP visualization of the PBMC30K^63^ dataset. Left: colored by cell type. Right: colored by batch. Prototypes are shown as dots with black edges. **(C)** Direct visualization of PBMC30K latent dimensions 13 and 14. **(D)** Evaluation of latent space disentanglement using the AtlasRGC^54^ dataset. Comparison of scIB metrics for the biological subspace (first-half latent space, **z**^1^) and the batch subspace (second-half latent space, **z**^2^). Metrics are grouped into biological conservation and batch correction categories. **(E)** UMAP visualization of AtlasRGC. **(F)** Direct visualization of AtlasRGC latent dimensions 13 and 14.

**Figure S2:**
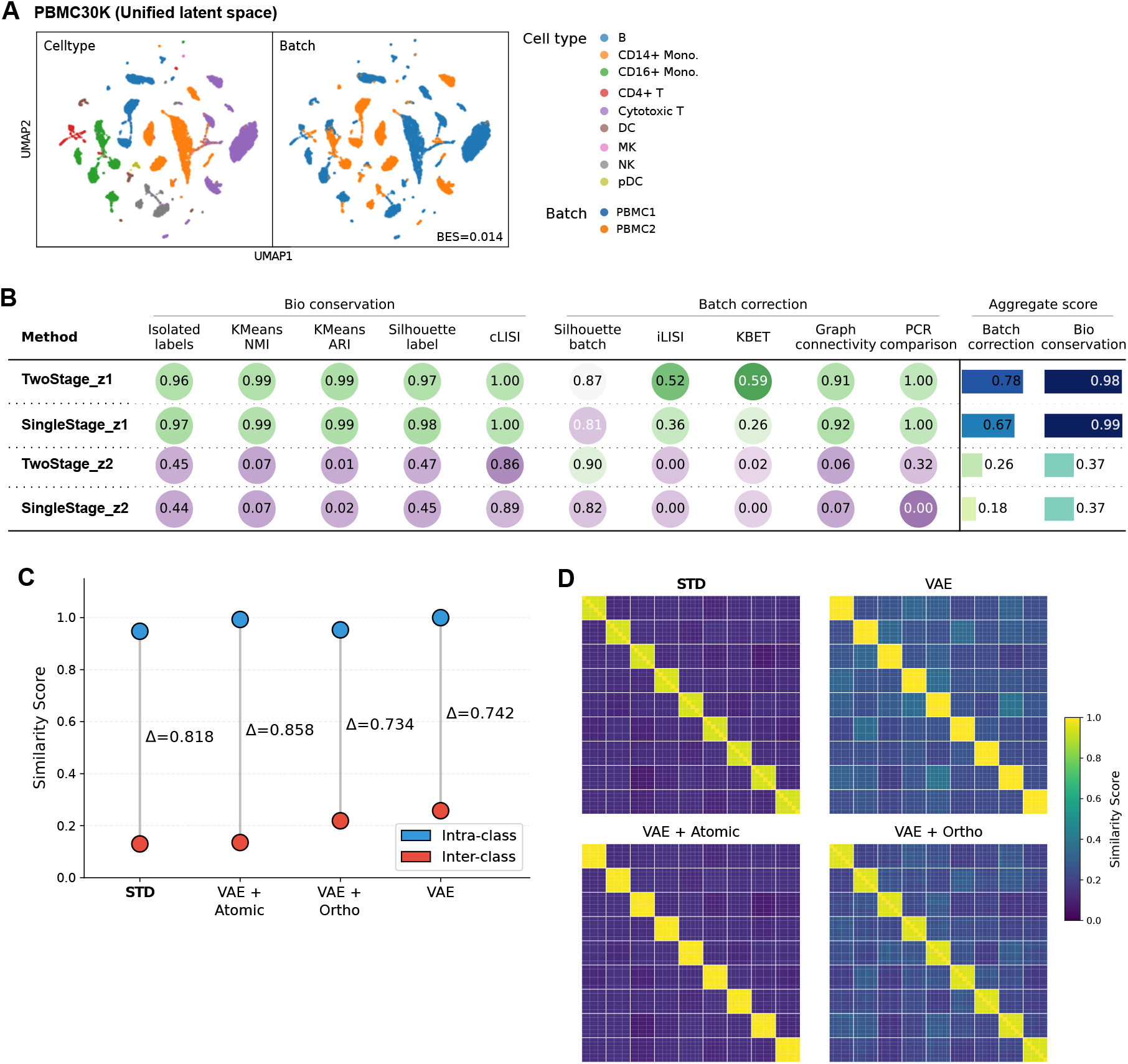
Ablation study of ProtoCloud model components, related to Figure 2-3. **(A)** UMAP visualization of ProtoCloud with unified latent space in the PBMC30K dataset, colored by cell type (left) and batch (right). **(B)** Comparison of scIB metrics between two-stage curriculum and single-stage end-to-end training. **(C)** Effect of loss components on prototype separation. Pairwise similarities are computed among prototypes, with intra-class similarity measured between prototypes of the same cell type and inter-class similarity between prototypes of different cell types. The four configurations are: standard ProtoCloud (STD), VAE with atomic loss only (VAE + Atomic), VAE with orthogonal loss only (VAE + Ortho), and VAE without both orthogonal and atomic losses (VAE). *Δ* indicates the difference between intra-class and inter-class similarity. **(D)** Heatmaps of pairwise prototype similarity for each loss configuration.

**Figure S3:**
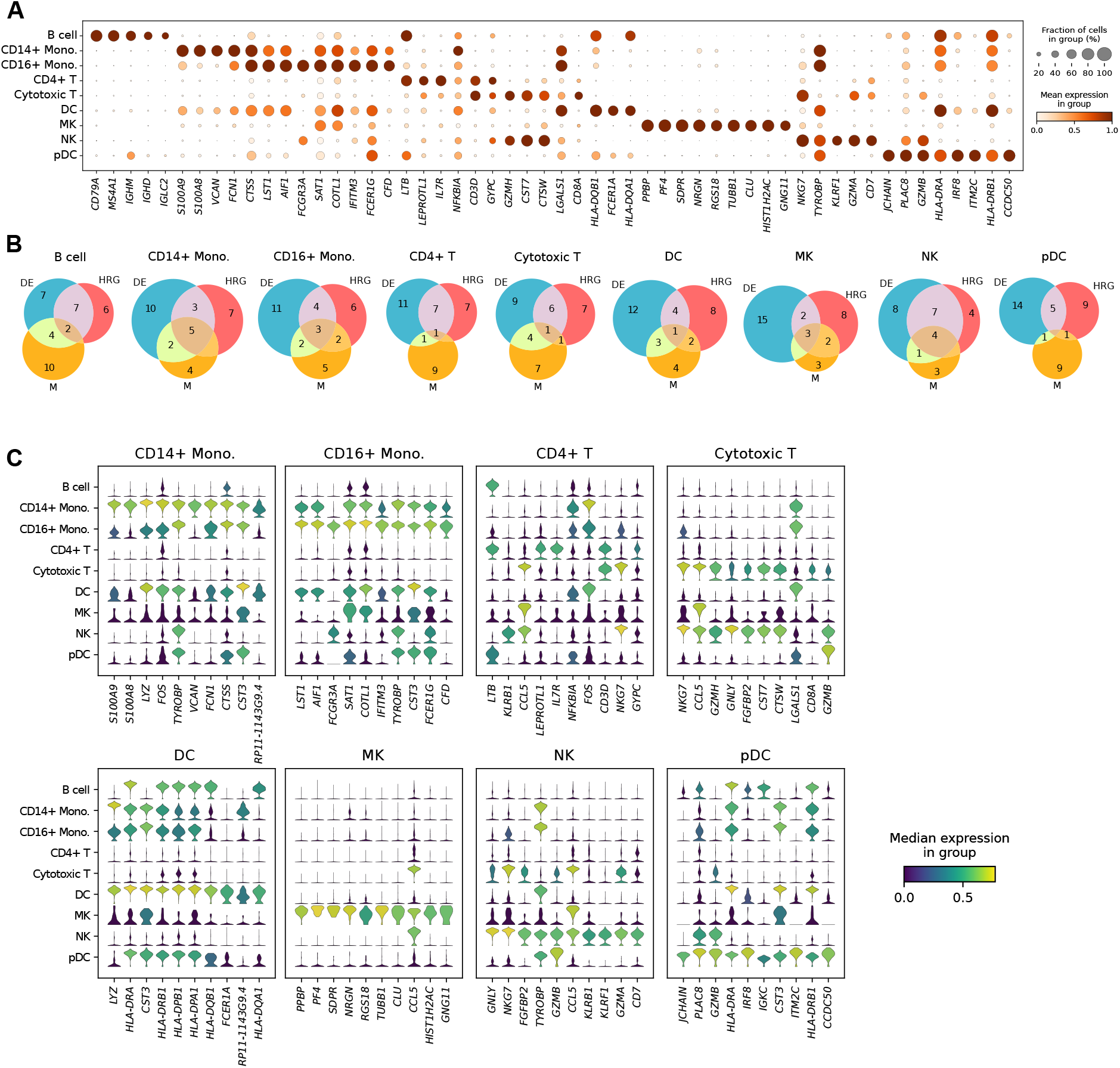
Highly relevant genes in the PBMC30K dataset^63^, related to Figure 3. **(A)** Dot plot of top ranked HRGs across cell types in PBMC30K. The gene set includes the top 10 HRGs for each cell type, where duplicated genes were removed to ensure a unique gene set. **(B)** Overlap of differentially expressed genes (DE), highly relevant genes (HRG), and canonical marker genes (M) for each cell type in PBMC30K. The overlap is shown with the top 20 DE genes, top 15 HRGs, and known markers (10-16 per type). **(C)** Violin plot of top HRGs for each cell type in PBMC30K. Each subplot highlights the distribution of the top ten HRG expression across different cell types.

**Figure S4:**
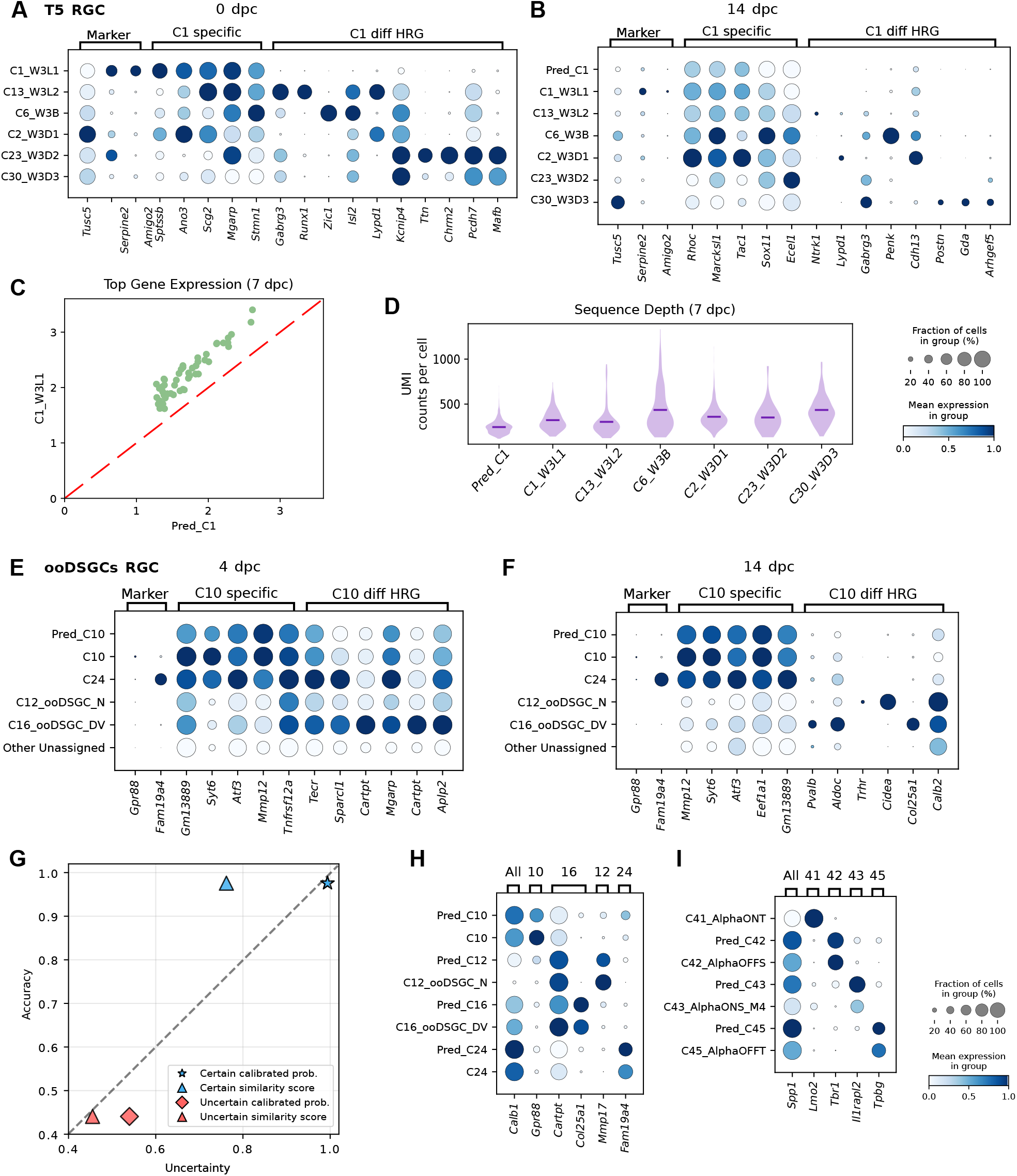
Characterization of predicted “unassigned” cells in comparison with reference RGC subtypes, related to Figure 5. **(A)-(B)** Comparison of HRG expression between W3-like C1 RGCs and other T5 RGCs, including unassigned cells predicted as C1 (Pred C1) cells. Dot plots showing the expression patterns of representative genes across different RGC subtypes at 0 (**A**) and 14 (**B**) days post-crush (dpc). Genes include known marker genes (*Tusc5, Serpine2*, and *Amigo2*), HRGs specific to C1 RGCs (C1 specific) and differentially ranked HRGs between C1 and other T5 RGCs. **(C)** The expression of the top highly expressed genes in predicted and reference C1 cells at 7 dpc. Points above the diagonal red dashed line indicate a systematic downregulation of these genes in predicted C1 cells compared to reference cells. **(D)** The distribution of sequencing depth across predicted C1 cells and reference T5 RGC subtypes at 7 dpc. **(E)-(F)** Comparison of HRG expression between C10 RGCs and other ooDSGCs RGCs, including unassigned cells predicted as C10 (Pred C10) cells. Dot plots showing the expression patterns of representative genes across different RGC subtypes at 4 dpc (**E**) and 14 dpc (**F**). **(G)** Reliability diagram of Patch-seq RGC certainty estimates. Both the model and the calibrator were trained on the AtlasRGC dataset and subsequently applied to Patch-seq RGC predictions. The plot compares uncertainty estimates derived from similarity scores and calibrated similarities for both certain and uncertain groups. **(H-I)** Validation of transferred annotations on Patch-seq RGC data. The dot plots compare the expression of marker genes between the most abundant predicted Patch-seq cells and their matched subtypes in AtlasRGC references for S2/S4 laminating RGCs (**H**) and *α* RGCs (**I**).

**Figure S5:**
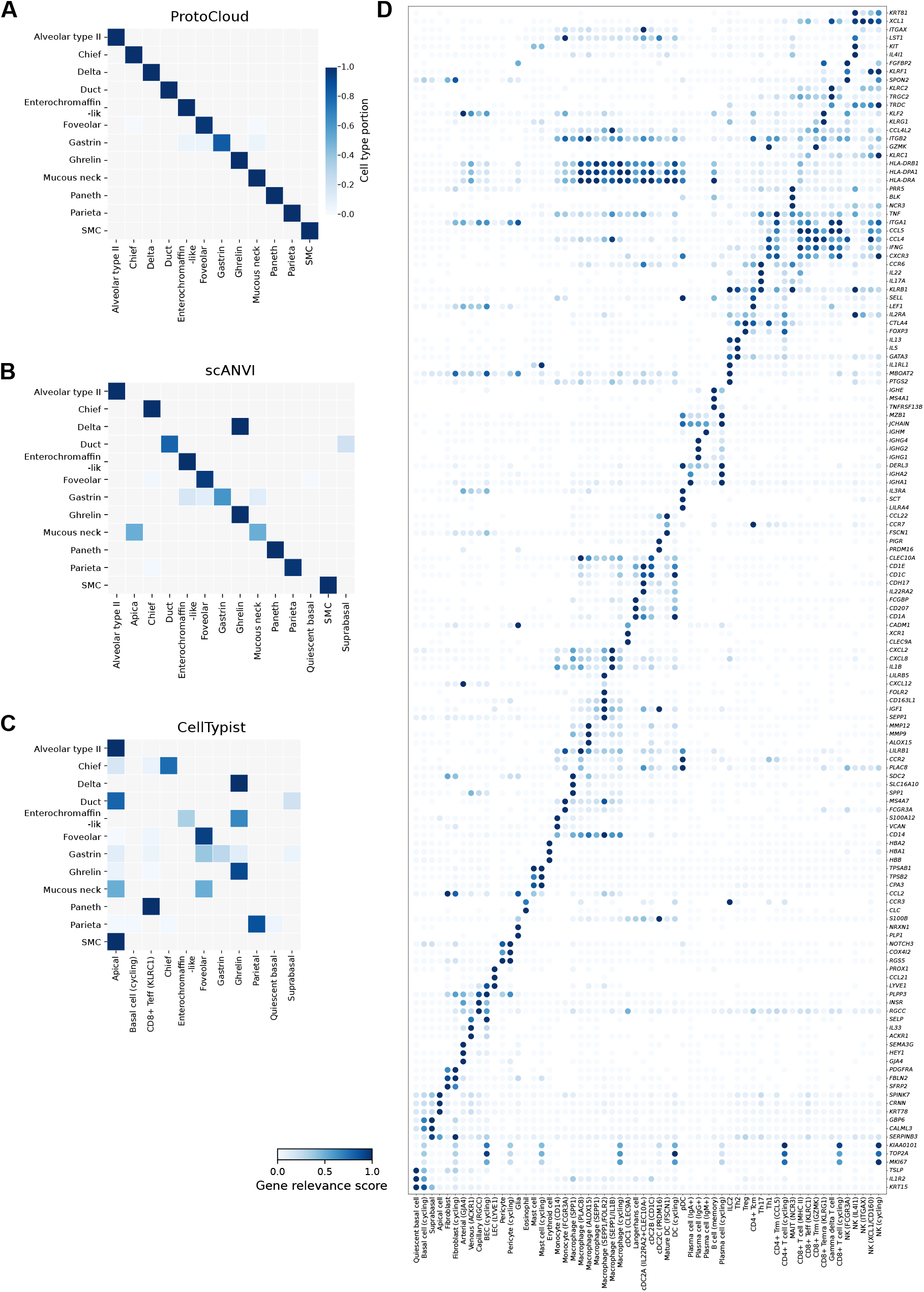
Model performance evaluation on AtlasEoE, related to Figure 6. **(A-C)** Confusion matrix of rare cell types in AtlasEoE^25^ across **(A)** ProtoCloud, **(B)** scANVI, and **(C)** CellTypist. **(D)** AtlasEoE marker genes relevance score dot plot. The dot plot illustrates the relevance scores of the provided marker genes (columns) for each of the 60 prevalent cell subsets (rows). This representation is adapted from the original study^25^, which used mean gene expression levels to represent marker gene–cell type relationships. The similarity in overall cell–gene relationships between the two approaches suggests that our relevance scores reliably capture the same biological trends as those reported in the original study.

**Figure S6:**
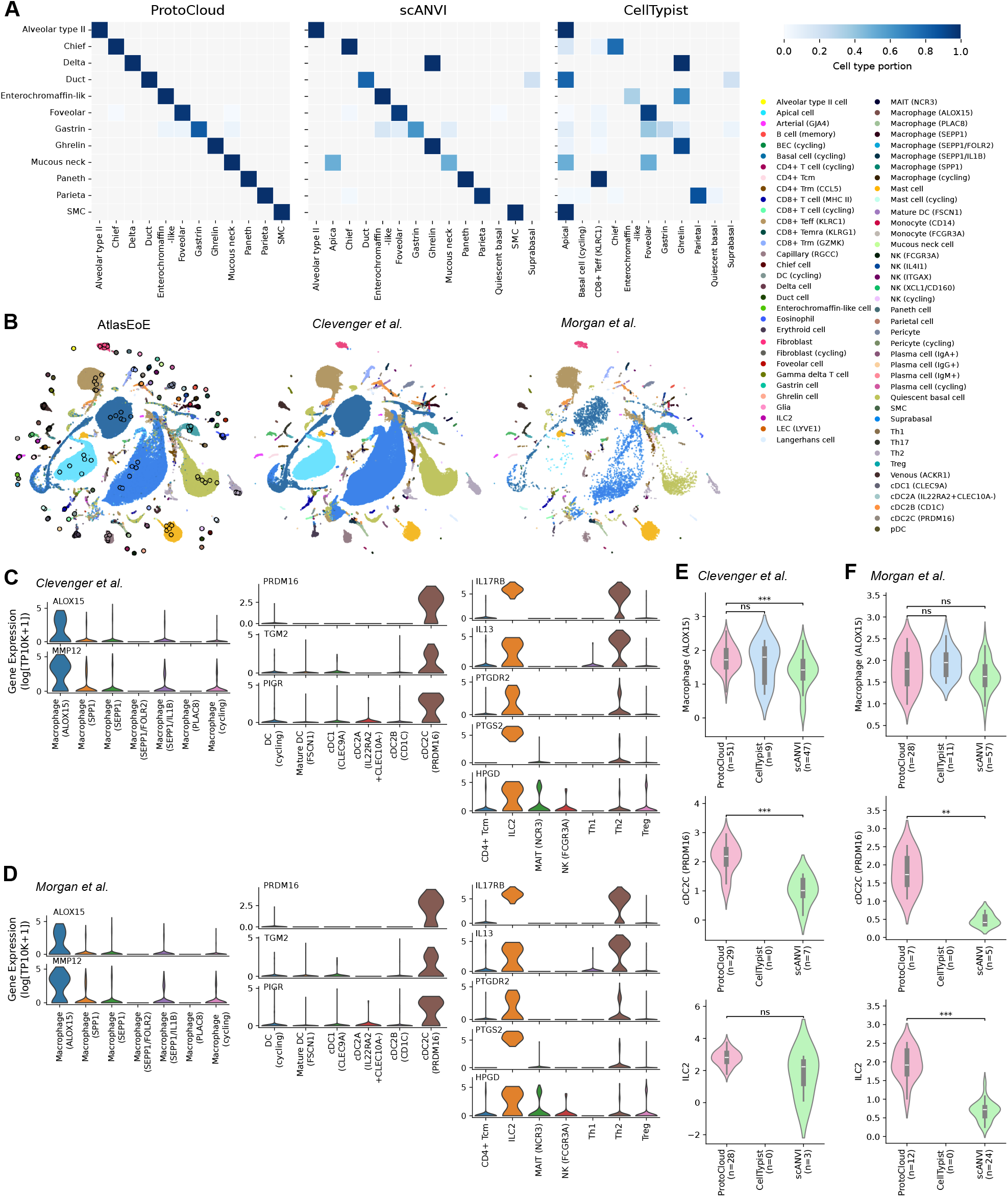
Model performance evaluation on applied esophageal tissue datasets, related to Figure 6. **(A)** UMAP visualization of the latent space embeddings for the AtlasEoE^25^ (left), Clevenger et al.^47^ (middle), and Morgan et al.^78^ (right) datasets. **(B)** Marker gene expression across cell types annotated by ProtoCloud in the Clevenger et al.^47^ dataset. Expression of marker genes *ALOX15* and *MMP12* in macrophage populations (left), *PRDM16, TGM2*, and *PIGR* expression across dendritic cell populations (middle), and ILC2 marker expression across several rare immune cell populations (right). **(C)** Corresponding marker expression patterns in the Morgan et al.^78^ dataset. **(D)** Specificity assessment of cell predictions in the Clevenger et al. dataset. Violin plots display the distribution of lineage-specific gene signature scores for *ALOX15* ^+^ macrophages (left), *PRDM16* ^+^ DCs (middle), and ILC2s (right). The plots compare the population identified by ProtoCloud (baseline) with non-overlapping cells identified exclusively by CellTypist or scANVI. Signature scores were calculated based on the top 30 marker genes derived from AtlasEoE (*n* indicates the number of cells; ns: non-significant, ** *p* ≤ 0.01, *** *p* ≤ 0.001). **(E)** Corresponding specificity assessment of cell predictions in the Morgan et al. dataset.

## Supplementary Tables

**Table S1:** Classification accuracy comparison between ProtoCloud and bench-mark methods across eight datasets, related to Figure 2. ProtoCloud results are shown as mean *±* standard error. Statistical significance is calculated using a paired t-test against ProtoCloud with Benjamini-Hochberg (FDR) correction (^∗^ *p* < 0.05, ^∗∗^ *p* < 0.01).

| Dataset | ProtoCloud | Seurat | scANVI | CellTypist | scPoli | TOSICA | SIMS | scGPT | scBERT |
| --- | --- | --- | --- | --- | --- | --- | --- | --- | --- |
| PBMC10K | **0.969 ± 0.003** | 0.977 ± 0.004* | 0.973 ± 0.004** | 0.952 ± 0.003** | 0.935 ± 0.010** | 0.967 ± 0.004 | 0.643 ± 0.039** | 0.856 ± 0.007** | 0.930 ± 0.034 |
| PBMC30K | **0.930 ± 0.005** | 0.942 ± 0.003* | 0.932 ± 0.003 | 0.917 ± 0.003** | 0.892 ± 0.004** | 0.912 ± 0.008* | 0.569 ± 0.032** | 0.804 ± 0.007** | 0.894 ± 0.025* |
| AtlasRGC | **0.979 ± 0.002** | 0.968 ± 0.002** | 0.976 ± 0.001* | 0.974 ± 0.002** | 0.943 ± 0.004** | 0.942 ± 0.010** | 0.287 ± 0.042** | 0.232 ± 0.004** | 0.973 ± 0.001** |
| TSCA_lung | **0.946 ± 0.001** | 0.927 ± 0.002** | 0.945 ± 0.002 | 0.927 ± 0.003** | 0.852 ± 0.007** | 0.915 ± 0.005** | 0.816 ± 0.023** | 0.624 ± 0.077** | 0.912 ± 0.019* |
| TSCA_oesophagus | **0.961 ± 0.002** | 0.951 ± 0.002** | 0.961 ± 0.001 | 0.938 ± 0.002** | 0.872 ± 0.005** | 0.942 ± 0.002** | 0.838 ± 0.121 | 0.811 ± 0.002** | 0.945 ± 0.005** |
| TSCA_spleen | **0.891 ± 0.005** | 0.857 ± 0.002** | 0.895 ± 0.003 | 0.865 ± 0.004** | 0.760 ± 0.005** | 0.830 ± 0.004** | 0.514 ± 0.064** | 0.608 ± 0.004** | / |
| AtlasEoE | **0.945 ± 0.002** | 0.916 ± 0.001** | 0.951 ± 0.001** | 0.925 ± 0.001** | 0.800 ± 0.007** | 0.927 ± 0.002** | 0.709 ± 0.095** | 0.584 ± 0.083** | / |
| ICA | **0.912 ± 0.002** | 0.879 ± 0.002** | 0.920 ± 0.001** | 0.891 ± 0.001** | 0.627 ± 0.003** | 0.844 ± 0.002** | 0.386 ± 0.029** | 0.537 ± 0.096** | / |

**Table S2:** Classification macro F1 comparison between ProtoCloud and bench-mark methods across eight datasets, related to Figure 2. ProtoCloud results are shown as mean *±* standard error. Statistical significance is calculated using a paired t-test against ProtoCloud with Benjamini-Hochberg (FDR) correction (^∗^ *p* < 0.05, ^∗∗^ *p* < 0.01).

| Dataset | ProtoCloud | Seurat | scANVI | CellTypist | scPoli | TOSICA | SIMS | scGPT | scBERT |
| --- | --- | --- | --- | --- | --- | --- | --- | --- | --- |
| PBMC10K | **0.930 ± 0.016** | 0.966 ± 0.009* | 0.938 ± 0.012 | 0.902 ± 0.022* | 0.902 ± 0.016* | 0.941 ± 0.006 | 0.264 ± 0.028** | 0.574 ± 0.013** | 0.772 ± 0.149 |
| PBMC30K | **0.917 ± 0.016** | 0.925 ± 0.011 | 0.912 ± 0.019 | 0.879 ± 0.028 | 0.847 ± 0.023** | 0.871 ± 0.040 | 0.278 ± 0.038** | 0.531 ± 0.037** | 0.726 ± 0.127 |
| AtlasRGC | **0.969 ± 0.002** | 0.933 ± 0.009** | 0.966 ± 0.002 | 0.954 ± 0.006** | 0.903 ± 0.005** | 0.929 ± 0.014** | 0.131 ± 0.039** | 0.091 ± 0.004** | 0.956 ± 0.005** |
| TSCA_lung | **0.924 ± 0.005** | 0.858 ± 0.004** | 0.911 ± 0.010* | 0.853 ± 0.006** | 0.757 ± 0.010** | 0.866 ± 0.014** | 0.453 ± 0.031** | 0.238 ± 0.048** | 0.804 ± 0.058** |
| TSCA_oesophagus | **0.955 ± 0.004** | 0.828 ± 0.017** | 0.942 ± 0.003** | 0.845 ± 0.019** | 0.676 ± 0.030** | 0.923 ± 0.006** | 0.299 ± 0.144** | 0.168 ± 0.002** | 0.805 ± 0.118* |
| TSCA_spleen | **0.862 ± 0.004** | 0.762 ± 0.005** | 0.856 ± 0.003* | 0.799 ± 0.005** | 0.653 ± 0.013** | 0.783 ± 0.007** | 0.226 ± 0.046** | 0.235 ± 0.006** | / |
| AtlasEoE | **0.877 ± 0.011** | 0.549 ± 0.015** | 0.841 ± 0.005** | 0.604 ± 0.009** | 0.397 ± 0.011** | 0.794 ± 0.007** | 0.100 ± 0.023** | 0.069 ± 0.016** | / |
| ICA | **0.888 ± 0.005** | 0.823 ± 0.009** | 0.889 ± 0.005 | 0.758 ± 0.009** | 0.524 ± 0.003** | 0.810 ± 0.005** | 0.079 ± 0.017** | 0.134 ± 0.036** | / |

**Table S3:** Rankings of learned HRGs for mature B cells from datasets of different organs, related to Figure 3. The three datasets used were from the lung, spleen, and esophagus of the Tissue Stability Cell Atlas. Because the esophagus dataset does not contain a mature B cell class, *CD27* ^+^ B cells were used instead.

| Gene | Dataset |  |  |
| --- | --- | --- | --- |
|  | Lung | Spleen | Esophagus |
| *EEF1A1* | 1 | 2 | 3 |
| *CD74* | 2 | 1 | 4 |
| *HLA-DRA* | 3 | 5 | 6 |
| *B2M* | 4 | 4 | 2 |
| *IGKC* | 5 | 10 | 5 |
| *MS4A1* | 6 | 30 | 41 |
| *TPT1* | 7 | 7 | 9 |
| *FTH1* | 8 | 22 | 15 |
| *CD79A* | 9 | 24 | 39 |
| *HLA-B* | 10 | 19 | 12 |

**Table S4:** Iterative training and evaluation on time-course RGC ONC dataset split by time post-crush, related to Figure 5. Only labeled cells are included in the evaluation. The ProtoCloud accuracies were computed from the final predictions after continue-training. CellTypist was initially trained on the control data (0 dpc) and used to predict labels for the data from 0.5 dpc. The confidently predicted 0.5 dpc cells (confidence score *>* 0.5) were then added to the training set, and the process was repeated for the data from each subsequent time point, progressively expanding the training set with predicted labels from earlier stages. scANVI followed the same incremental training procedure, but included all predicted cells during continue training as the model does not provide confidence estimates.

| DPC | Cells | ProtoCloud |  |  |  | CellTypist |  |  |  | scANVI |
| --- | --- | --- | --- | --- | --- | --- | --- | --- | --- | --- |
|  |  | Acc. | Certain (%) | Certain Acc. | Amb. Acc. | Acc. | Certain (%) | Certain Acc. | Amb. Acc. | Acc. |
| 0 (ctrl) | 12062 | 0.949 | 73.1% | 0.992 | 0.834 | **0.955** | 72.3% | 0.985 | 0.877 | 0.950 |
| 0.5 | 13619 | 0.938 | 78.4% | 0.988 | 0.759 | **0.944** | 57.2% | 0.988 | 0.886 | 0.943 |
| 1 | 12478 | **0.921** | 86.6% | 0.964 | 0.641 | 0.906 | 38.3% | 0.989 | 0.855 | 0.904 |
| 2 | 10695 | **0.863** | 83.7% | 0.929 | 0.524 | 0.837 | 15.1% | 0.973 | 0.812 | 0.838 |
| 4 | 10599 | **0.656** | 69.1% | 0.766 | 0.410 | 0.483 | 2.8% | 0.626 | 0.479 | 0.533 |
| 7 | 8700 | **0.627** | 74.5% | 0.717 | 0.365 | 0.399 | 4.8% | 0.581 | 0.390 | 0.506 |
| 14 | 8456 | **0.685** | 82.6% | 0.748 | 0.387 | 0.396 | 4.7% | 0.802 | 0.376 | 0.616 |

**Table S5:** Percentage of confidently labeled previously unknown cells at varying DPC concentrations, related to Figure 5. Shown are the number of total cells, previously “unknown” cells across different time points, and the percentage of these previously unknown cells that were confidently identified by ProtoCloud and CellTypist. Over the time course, approximately 20% to 70% of previously unknown cells were confidently assigned to a cell type by ProtoCloud.

| DPC | Cells | “Unknown” cells | Certain “unknown” (%) |  |
| --- | --- | --- | --- | --- |
|  |  |  | ProtoCloud | CellTypist |
| 0 (ctrl) | 12062 | 0 | / | / |
| 0.5 | 13619 | 157 | 22.3% | 56.6% |
| 1 | 12478 | 370 | 42.2% | 37.2% |
| 2 | 10695 | 501 | 48.5% | 14.4% |
| 4 | 10599 | 2522 | 62.5% | 4.1% |
| 7 | 8700 | 2178 | 72.4% | 8.2% |
| 14 | 8456 | 1455 | 73.6% | 4.8% |

**Table S6:**
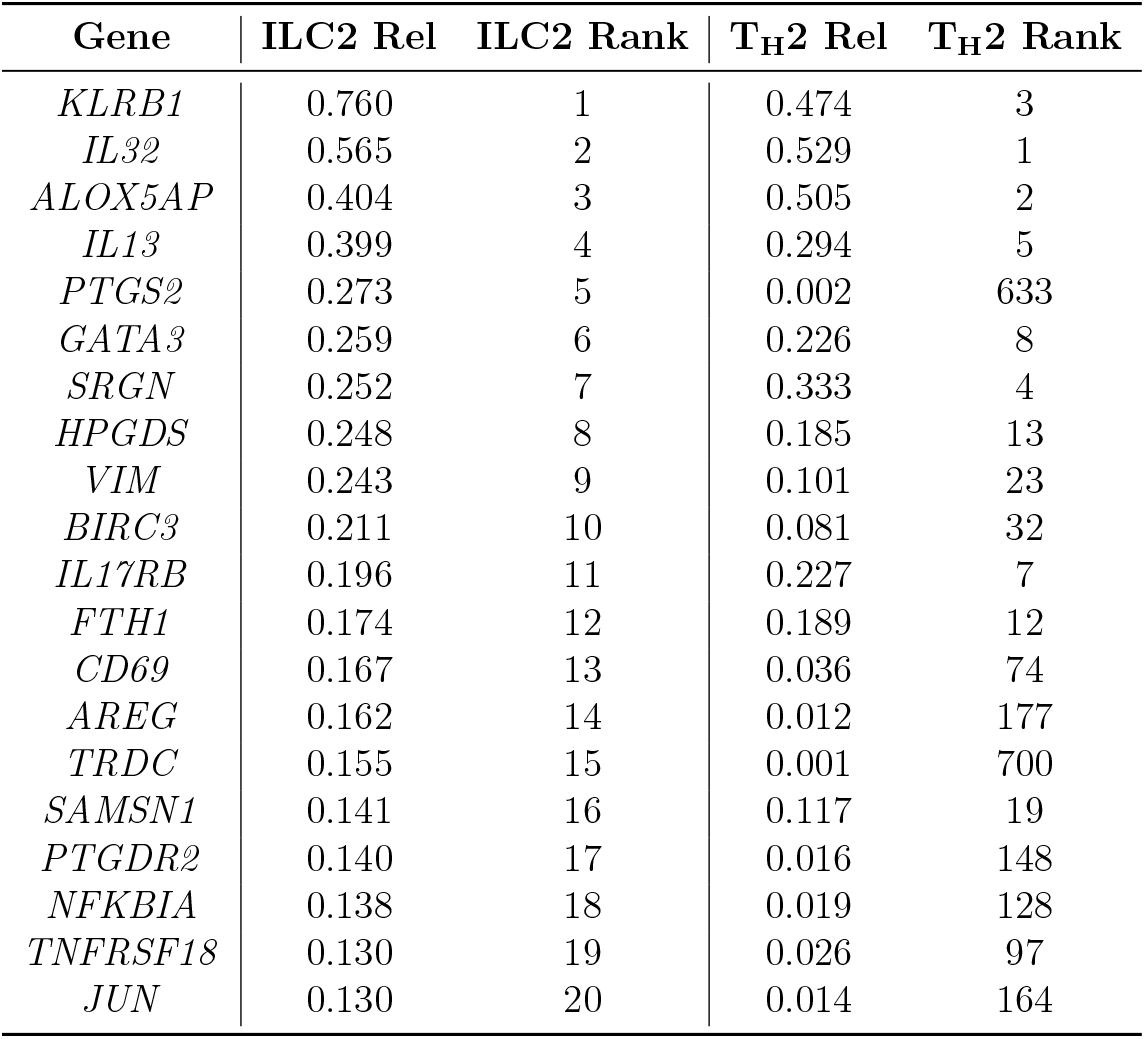
Top ILC2 highly relevant genes in AtlasEoE, related to Figure 6. The top ILC2 HRGs, along with their relevance and ranks in ILC2s and T_H_2 cells, highlight the upregulation of the prostaglandin-related genes *PTGS2* and *PTGDR2*, and the epidermal growth factor family gene *AREG*.

**Table S7:** Cell type identification among ProtoCloud, CellTypist, and scANVI across two applied EoE datasets, related to Figure 6. While all models identify common populations, ProtoCloud demonstrates a higher subtype resolution compared to CellTypist. CellTypist shows reduced sensitivity in detecting rare populations, failing to identify ILC2s in the Clevenger dataset and cycling DCs in the Morgan dataset. Although scANVI shows high sensitivity for these rare cell states, some predictions may be of low quality (**Figure** S6).

| Cell type | Clevenger et al. |  |  | Morgan et al. |  |  |
| --- | --- | --- | --- | --- | --- | --- |
|  | ProtoCloud | CellTypist | scANVI | ProtoCloud | CellTypist | scANVI |
| Macrophage (*SEPP1*) | 905 | 1607 | 1061 | 25 | 86 | 67 |
| Macrophage (*SEPP1/IL1B*) | 662 | 357 | 391 | 355 | 321 | 294 |
| Macrophage (*cycling*) | 511 | 285 | 311 | 22 | 11 | 13 |
| Macrophage (*PLAC8*) | 283 | 147 | 501 | 90 | 19 | 53 |
| Macrophage (*ALOX15*) | 51 | 19 | 81 | 28 | 20 | 74 |
| Macrophage (*SPP1*) | 3 | 1 | 7 | 1 | 10 | 67 |
| Macrophage (*SEPP1/FOLR2*) | 4 | 1 | 7 | 2 | 1 | 6 |
| Treg | 1679 | 1967 | 1983 | 609 | 562 | 821 |
| T<sub>H</sub>2 | 283 | 266 | 283 | 207 | 202 | 367 |
| *CD4* + Tcm | 429 | 55 | 171 | 176 | 90 | 75 |
| MAIT (*NCR3*) | 209 | 8 | 256 | 45 | 11 | 21 |
| T<sub>H</sub>1 | 65 | 2 | 55 | 14 | 18 | 7 |
| NK (*FCGR3A*) | 45 | 5 | 44 | 9 | 1 | 7 |
| ILC2 | 28 | **0** | 26 | 12 | 1 | 30 |
| cDC2B (*CD1C*) | 1980 | 1450 | 2076 | 179 | 151 | 180 |
| Mature DC (*FSCN1*) | 563 | 324 | 378 | 28 | 49 | 26 |
| cDC1 (*CLEC9A*) | 201 | 141 | 239 | 31 | 24 | 34 |
| DC (*cycling*) | 240 | 64 | 266 | 3 | **0** | 3 |
| cDC2A (*IL22RA2*+*CLEC10A*<sup>-</sup>) | 50 | 7 | 43 | 1 | 1 | 2 |
| cDC2C (*PRDM16*) | 29 | 16 | 23 | 7 | 3 | 10 |

## References

1. Klein, A. M., Mazutis, L., Akartuna, I., Tallapragada, N., Veres, A., Li, V., Peshkin, L., Weitz, D. A., and Kirschner, M. W. (2015). Droplet barcoding for single-cell transcriptomics applied to embryonic stem cells. Cell 161, 1187–1201.

2. Macosko, E. Z., Basu, A., Satija, R., Nemesh, J., Shekhar, K., Goldman, M., Tirosh, I., Bialas, A. R., Kamitaki, N., Martersteck, E. M., et al. (2015). Highly parallel genomewide expression profiling of individual cells using nanoliter droplets. Cell 161, 1202–1214.

3. Zheng, G. X., Terry, J. M., Belgrader, P., Ryvkin, P., Bent, Z. W., Wilson, R., Ziraldo, S. B., Wheeler, T. D., McDermott, G. P., Zhu, J., et al. (2017). Massively parallel digital transcriptional profiling of single cells. Nature Communications 8, 14049.

4. Cao, J., Packer, J. S., Ramani, V., Cusanovich, D. As, Huynh, C., Daza, R., Qiu, X., Lee, C., Furlan, S. N., Steemers, F. J., et al. (2017). Comprehensive single-cell transcriptional profiling of a multicellular organism. Science 357, 661–667.

5. Rosenberg, A. B., Roco, C. M., Muscat, R. A., Kuchina, A., Sample, P., Yao, Z., Graybuck, L. T., Peeler, D. J., Mukherjee, S., Chen, W., et al. (2018). Single-cell profiling of the developing mouse brain and spinal cord with split-pool barcoding. Science 360, 176–182.

6. Gierahn, T. M., Wadsworth, M. H., Hughes, T. K., Bryson, B. D., Butler, A., Satija, R., Fortune, S., Love, J. C., and Shalek, A. K. (2017). Seq-Well: portable, low-cost RNA sequencing of single cells at high throughput. Nature Methods 14, 395–398.

7. Han, X., Wang, R., Zhou, Y., Fei, L., Sun, H., Lai, S., Saadatpour, A., Zhou, Z., Chen, H., Ye, F., et al. (2018). Mapping the mouse cell atlas by microwell-seq. Cell 172, 1091–1107.

8. Domínguez Conde, C., Xu, C., Jarvis, L., Rainbow, D., Wells, S., Gomes, T., Howlett, S., Suchanek, O., Polanski, K., King, H., et al. (2022a). Cross-tissue immune cell analysis reveals tissue-specific features in humans. Science 376, eabl5197. 10.1126/science.abl5197.

9. Travaglini, K. J., Nabhan, A. N., Penland, L., Sinha, R., Gillich, A., Sit, R. V., Chang, S., Conley, S. D., Mori, Y., Seita, J., et al. (2020). A molecular cell atlas of the human lung from single-cell RNA sequencing. Nature 587, 619–625.

10. Wagner, J., Rapsomaniki, M. A., Chevrier, S., Anzeneder, T., Langwieder, C., Dykgers, A., Rees, M., Ramaswamy, A., Muenst, S., Soysal, S. D., et al. (2019). A single-cell atlas of the tumor and immune ecosystem of human breast cancer. Cell 177, 1330–1345.

11. Regev, A., Teichmann, S. A., Lander, E. S., Amit, I., Benoist, C., Birney, E., Bodenmiller, B., Campbell, P., Carninci, P., Clatworthy, M., et al. (2017). The human cell atlas. eLife 6, e27041.

12. Smillie, C. S., Biton, M., Ordovas-Montanes, J., Sullivan, K. M., Burgin, G., Graham, D. B., Herbst, R. H., Rogel, N., Slyper, M., Waldman, J., et al. (2019). Intra-and inter-cellular rewiring of the human colon during ulcerative colitis. Cell 178, 714–730.

13. Xu, H., Ding, J., Porter, C. B., Wallrapp, A., Tabaka, M., Ma, S., Fu, S., Guo, X., Riesenfeld, S. J., Su, C., et al. (2019). Transcriptional atlas of intestinal immune cells reveals that neuropeptide α-CGRP modulates group 2 innate lymphoid cell responses. Immunity 51, 696–708.

14. Efremova, M., Vento-Tormo, M., Teichmann, S. A., and Vento-Tormo, R. (2020). Cell-PhoneDB: inferring cell–cell communication from combined expression of multi-subunit ligand–receptor complexes. Nature Protocols 15, 1484–1506.

15. Aizarani, N., Saviano, A., Sagar, n., Mailly, L., Durand, S., Herman, J. S., Pessaux, P., Baumert, T. F., and Grün, D. (2019). A human liver cell atlas reveals heterogeneity and epithelial progenitors. Nature 572, 199–204.

16. Plasschaert, L. W., Žilionis, R., Choo-Wing, R., Savova, V., Knehr, J., Roma, G., Klein, M., and Jaffe, A. B. (2018). A single-cell atlas of the airway epithelium reveals the CFTR-rich pulmonary ionocyte. Nature 560, 377–381.

17. Gavish, A., Tyler, M., Greenwald, A. C., Hoefflin, R., Simkin, D., Tschernichovsky, R., Galili Darnell, N., Somech, E., Barbolin, C., Antman, T., et al. (2023). Hallmarks of transcriptional intratumour heterogeneity across a thousand tumours. Nature 618, 598–606.

18. Karaiskos, N., Wahle, P., Alles, J., Boltengagen, A., Ayoub, S., Kipar, C., Kocks, C., Rajewsky, N., and Zinzen, R. P. (2017). The Drosophila embryo at single-cell transcriptome resolution. Science 358, 194–199.

19. Wagner, D. E., Weinreb, C., Collins, Z. M., Briggs, J. A., Megason, S. G., and Klein, M. (2018). Single-cell mapping of gene expression landscapes and lineage in the zebrafish embryo. Science 360, 981–987.

20. The Tabula Muris Consortium (2020). A single-cell transcriptomic atlas characterizes ageing tissues in the mouse. Nature 583, 590–595.

21. He, S., Wang, L.-H., Liu, Y., Li, Y.-Q., Chen, H.-T., Xu, J.-H., Peng, W., Lin, G.-W., Wei, P.-P., Li, B., et al. (2020). Single-cell transcriptome profiling of an adult human cell atlas of 15 major organs. Genome Biology 21, 294.

22. Jin, K., Yao, Z., Velthoven, C. T. van, Kaplan, E. S., Glattfelder, K., Barlow, S. T., Boyer, G., Carey, D., Casper, T., Chakka, A. B., et al. (2025). Brain-wide cell-type-specific transcriptomic signatures of healthy ageing in mice. Nature 638, 182–196.

23. Vázquez-García, I., Uhlitz, F., Ceglia, N., Lim, J. L., Wu, M., Mohibullah, N., Niyazov, J., Ruiz, A. E. B., Boehm, K. M., Bojilova, V., et al. (2022). Ovarian cancer mutational processes drive site-specific immune evasion. Nature 612, 778–786.

24. Alladina, J., Smith, N. P., Kooistra, T., Slowikowski, K., Kernin, I. J., Deguine, J., Keen, H. L., Manakongtreecheep, K., Tantivit, J., Rahimi, R. A., et al. (2023). A human model of asthma exacerbation reveals transcriptional programs and cell circuits specific to allergic asthma. Science Immunology 8, eabq6352.

25. Ding, J., Garber, J. J., Uchida, A., Lefkovith, A., Carter, G. T., Vimalathas, P., Canha, L., Dougan, M., Staller, K., Yarze, J., et al. (2024). An esophagus cell atlas reveals dynamic rewiring during active eosinophilic esophagitis and remission. Nature Communications 15, 3344. 10.1038/s41467-024-47647-0.

26. Xu, C., Lopez, R., Mehlman, E., Regier, J., Jordan, M. I., and Yosef, N. (2021). Probabilistic harmonization and annotation of single-cell transcriptomics data with deep generative models. Molecular Systems Biology 17, e9620. 10.15252/msb.20209620.

27. Hao, Y., Hao, S., Andersen-Nissen, E., Mauck, W. M., Zheng, S., Butler, A., Lee, M. J., Wilk, A. J., Darby, C., Zager, M., et al. (2021). Integrated analysis of multimodal singlecell data. Cell 184, 3573–3587. 10.1016/j.cell.2021.04.048.

28. Domínguez Conde, C., Xu, C., Jarvis, L. B., Rainbow, D. B., Wells, S. B., Gomes, T., Howlett, S. K., Suchanek, O., Polanski, K., King, H. W., et al. (2022b). Cross-tissue immune cell analysis reveals tissue-specific features in humans. Science 376, eabl5197. 10.1126/science.abl5197.

29. Ergen, C., Xing, G., Xu, C., Kim, M., Jayasuriya, M., McGeever, E., Oliveira Pisco, A., Streets, A., and Yosef, N. (2024). Consensus prediction of cell type labels in single-cell data with popV. Nature Genetics 56, 2731–2738.

30. Satija, R., Farrell, J. A., Gennert, D., Schier, A. F., and Regev, A. (2015). Spatial reconstruction of single-cell gene expression data. Nature biotechnology 33, 495–502.

31. Ianevski, A., Giri, A. K., and Aittokallio, T. (2022). Fully-automated and ultra-fast celltype identification using specific marker combinations from single-cell transcriptomic data. Nature communications 13, 1246.

32. Ding, J., Condon, A., and Shah, S. P. (2018). Interpretable dimensionality reduction of single cell transcriptome data with deep generative models. Nature Communications 9, 2002.

33. Lopez, R., Regier, J., Cole, M. B., Jordan, M. I., and Yosef, N. (2018). Deep generative modeling for single-cell transcriptomics. Nature Methods 15, 1053–1058. 10.1038/s41592-018-0229-2.

34. Amodio, M., Van Dijk, D., Srinivasan, K., Chen, W. S., Mohsen, H., Moon, K. R., Campbell, A., Zhao, Y., Wang, X., Venkataswamy, M., et al. (2019). Exploring single-cell data with deep multitasking neural networks. Nature Methods 16, 1139–1145.

35. Eraslan, G., Simon, L. M., Mircea, M., Mueller, N. S., and Theis, F. J. (2019). Single-cell RNA-seq denoising using a deep count autoencoder. Nature Communications 10, 390.

36. Li, X., Wang, K., Lyu, Y., Pan, H., Zhang, J., Stambolian, D., Susztak, K., Reilly, M. P., Hu, G., and Li, M. (2020). Deep learning enables accurate clustering with batch effect removal in single-cell RNA-seq analysis. Nature Communications 11, 2338.

37. Ding, J. and Regev, A. (2021). Deep generative model embedding of single-cell RNA-Seq profiles on hyperspheres and hyperbolic spaces. Nature Communications 12, 2554.

38. Yang, F., Wang, W., Wang, F., Fang, Y., Tang, D., Huang, J., Lu, H., and Yao, J. (2022). scBERT as a large-scale pretrained deep language model for cell type annotation of single-cell RNA-seq data. Nature Machine Intelligence 4, 852–866.

39. Wen, H., Tang, W., Dai, X., Ding, J., Jin, W., Xie, Y., and Tang, J. (2023). CellPLM: pre-training of cell language model beyond single cells. bioRxiv, 2023–10. 10.1101/2023.10.03.560734.

40. Cui, H., Wang, C., Maan, H., Pang, K., Luo, F., Duan, N., and Wang, B. (2024). scGPT: toward building a foundation model for single-cell multi-omics using generative AI. Nature Methods 21, 1470–1480.

41. Karin, J., Mintz, R., Raveh, B., and Nitzan, M. (2024). Interpreting single-cell and spatial omics data using deep neural network training dynamics. Nature Computational Science 4, 941–954.

42. Arrieta, A. B., Díaz-Rodríguez, N., Del Ser, J., Bennetot, A., Tabik, S., Barbado, A., García, S., Gil-López, S., Molina, D., Benjamins, R., et al. (2020). Explainable Artificial Intelligence (XAI): Concepts, taxonomies, opportunities and challenges toward responsible AI. Information Fusion 58, 82–115.

43. Alvarez Melis, D. and Jaakkola, T. (2018). “Towards Robust Interpretability with Self-Explaining Neural Networks”. Advances in Neural Information Processing Systems. Vol. 31.

44. Johansen, N. and Quon, G. (2019). scAlign: a tool for alignment, integration, and rare cell identification from scRNA-seq data. Genome Biology 20, 166.

45. Falcone, F. H., Haas, H., and Gibbs, B. F. (2000). The human basophil: a new appreciation of its role in immune responses. Blood 96, 4028–4038.

46. Rochman, M., Wen, T., Kotliar, M., Dexheimer, P. J., Morgenstern, N. B.-B., Cald-well, J. M., Lim, H.-W., and Rothenberg, M. E. (2022). Single-cell RNA-Seq of human esophageal epithelium in homeostasis and allergic inflammation. JCI Insight 7.

47. Clevenger, M. H., Karami, A. L., Carlson, D. A., Kahrilas, P. J., Gonsalves, N., Pandolfino, J. E., Winter, D. R., Whelan, K. A., and Tétreault, M.-P. (2023). Suprabasal cells retain progenitor cell identity programs in eosinophilic esophagitis–driven basal cell hyperplasia. JCI Insight 8. 10.1172/jci.insight.171765.

48. Morgenstern, N. B.-B., Rochman, M., Kotliar, M., Dunn, J. L., Mack, L., Besse, J., Natale, M. A., Klingler, A. M., Felton, J. M., Caldwell, J. M., et al. (2024). Single-cell RNA Sequencing of Human Eosinophils in Allergic Inflammation in the Esophagus. Journal of Allergy and Clinical Immunology 154, 974–987.

49. Mereu, E., Lafzi, A., Moutinho, C., Ziegenhain, C., McCarthy, D. J., Aĺvarez-Varela, A., Batlle, E., Sagar, n., Gruen, D., Lau, J. K., et al. (2020). Benchmarking single-cell RNA-sequencing protocols for cell atlas projects. Nature Biotechnology 38, 747–755.

50. Lähnemann, D., Köster, J., Szczurek, E., McCarthy, D. J., Hicks, S. C., Robinson, M. D., Vallejos, C. A., Campbell, K. R., Beerenwinkel, N., Mahfouz, A., et al. (2020). Eleven grand challenges in single-cell data science. Genome Biology 21, 31.

51. Korsunsky, I., Millard, N., Fan, J., Slowikowski, K., Zhang, F., Wei, K., Baglaenko, Y., Brenner, M., Loh, P.-r., and Raychaudhuri, S. (2019). Fast, sensitive and accurate integration of single-cell data with Harmony. Nature Methods 16, 1289–1296.

52. Chen, S., Regev, A., Condon, A., and Ding, J. (2026). CellUntangler: separating distinct biological signals in single-cell data with deep generative models. Cell Genomics 6.

53. Wang, Z., Yeo, G. H., Sherwood, R., and Gifford, D. (2019). “Disentangled representations of cellular identity”. Research in Computational Molecular Biology. Springer International Publishing, 256–271. ISBN: 978-3-030-17083-7.

54. Tran, N. M., Shekhar, K., Whitney, I. E., Jacobi, A., Benhar, I., Hong, G., Yan, W., Adiconis, X., Arnold, M. E., Lee, J. M., et al. (2019). Single-cell profiles of retinal ganglion cells differing in resilience to injury reveal neuroprotective genes. Neuron 104, 1039–1055. 10.1016/j.neuron.2019.11.006.

55. Kingma, D. P. (2014). “Auto-encoding variational bayes”. 2nd International Conference on Learning Representations.

56. Rezende, D. J., Mohamed, S., and Wierstra, D. (2014). “Stochastic backpropagation and approximate inference in deep generative models”. International Conference on Machine Learning. Vol. 32. PMLR, 1278–1286.

57. Bach, S., Binder, A., Montavon, G., Klauschen, F., Müller, K.-R., and Samek, W. (2015). On pixel-wise explanations for non-linear classifier decisions by layer-wise rele-vance propagation. PLOS ONE 10, 1–46.

58. Gautam, S., Höhne, M. M.-C., Hansen, S., Jenssen, R., and Kampffmeyer, M. (2023). This looks More Like that: Enhancing Self-Explaining Models by Prototypical Rele-vance Propagation. Pattern Recognition 136, 109172. 10.1016/j.patcog.2022.109172.

59. Gautam, S., Boubekki, A., Hansen, S., Salahuddin, S., Jenssen, R., Höhne, M., and Kampffmeyer, M. (2022). ProtoVAE: A trustworthy self-explainable prototypical variational model. Advances in Neural Information Processing Systems 35, 17940–17952.

60. Chen, J., Xu, H., Tao, W., Chen, Z., Zhao, Y., and Han, J.-D. J. (2023). Transformer for one stop interpretable cell type annotation. Nature Communications 14, 223.

61. De Donno, C., Hediyeh-Zadeh, S., Moinfar, A. A., Wagenstetter, M., Zappia, L., Lotfollahi, M., and Theis, F. J. (2023). Population-level integration of single-cell datasets enables multi-scale analysis across samples. Nature Methods 20, 1683–1692.

62. Gonzalez-Ferrer, J., Lehrer, J., O’Farrell, A., Paten, B., Teodorescu, M., Haussler, D., Jonsson, V. D., and Mostajo-Radji, M. A. (2024). SIMS: A deep-learning label transfer tool for single-cell RNA sequencing analysis. Cell Genomics 4.

63. Ding, J., Adiconis, X., Simmons, S. K., Kowalczyk, M. S., Hession, C. C., Marjanovic, N. D., Hughes, T. K., Wadsworth, M. H., Burks, T., Nguyen, L. T., et al. (2020). Systematic comparison of single-cell and single-nucleus RNA-sequencing methods. Nature Biotechnology 38, 737–746.

64. Madissoon, E., Wilbrey-Clark, A., Miragaia, R. J., Saeb-Parsy, K., Mahbubani, K. T., Georgakopoulos, N., Harding, P., Polanski, K., Huang, N., Nowicki-Osuch, K., et al. (2019). scRNA-seq assessment of the human lung, spleen, and esophagus tissue stability after cold preservation. Genome Biology 21. 10.1186/s13059-019-1906-x.

65. McHugh, M. L. (2012). Interrater reliability: the kappa statistic. Biochemia medica 22, 276–282.

66. Rautenstrauch, P. and Ohler, U. (2025). Shortcomings of silhouette in single-cell integration benchmarking. Nature Biotechnology, 1–5.

67. McInnes, L., Healy, J., and Melville, J. (2018). UMAP: Uniform manifold approximation and projection for dimension reduction. arXiv preprint arXiv:1802.03426.

68. Zhang, X., Lan, Y., Xu, J., Quan, F., Zhao, E., Deng, C., Luo, T., Xu, L., Liao, G., Yan, M., et al. (2019a). CellMarker: a manually curated resource of cell markers in human and mouse. Nucleic Acids Research 47, D721–D728.

69. Zhang, Z., Luo, D., Zhong, X., Choi, J. H., Ma, Y., Wang, S., Mahrt, E., Guo, W., Stawiski, E. W., Modrusan, Z., et al. (2019b). SCINA: a semi-supervised subtyping algorithm of single cells and bulk samples. Genes 10, 531.

70. Zhang, A. W., O’Flanagan, C., Chavez, E. A., Lim, J. L., Ceglia, N., McPherson, A., Wiens, M., Walters, P., Chan, T., Hewitson, B., et al. (2019c). Probabilistic cell-type assignment of single-cell RNA-seq for tumor microenvironment profiling. Nature Methods 16, 1007–1015.

71. Tirosh, I., Izar, B., Prakadan, S. M., Wadsworth, M. H., Treacy, D., Trombetta, J. J., Rotem, A., Rodman, C., Lian, C., Murphy, G., et al. (2016). Dissecting the multicellular ecosystem of metastatic melanoma by single-cell RNA-seq. Science 352, 189–196.

72. Yanai, I., Benjamin, H., Shmoish, M., Chalifa-Caspi, V., Shklar, M., Ophir, R., Bar-Even, A., Horn-Saban, S., Safran, M., Domany, E., et al. (2005). Genome-wide midrange transcription profiles reveal expression level relationships in human tissue specification. Bioinformatics 21, 650–659.

73. Glenn, W. B. et al. (1950). Verification of forecasts expressed in terms of probability. Monthly weather review 78, 1–3.

74. Nixon, J., Dusenberry, M. W., Zhang, L., Jerfel, G., and Tran, D. (2019). “Measuring Calibration in Deep Learning”. Proceedings of the IEEE/CVF Conference on Computer Vision and Pattern Recognition (CVPR) Workshops.

75. Zadrozny, B. and Elkan, C. (2002). “Transforming classifier scores into accurate multiclass probability estimates”. Proceedings of the eighth ACM SIGKDD international conference on Knowledge discovery and data mining, 694–699.

76. Huang, W., Xu, Q., Su, J., Tang, L., Hao, Z.-Z., Xu, C., Liu, R., Shen, Y., Sang, X., Xu, N., et al. (2022). Linking transcriptomes with morphological and functional phenotypes in mammalian retinal ganglion cells. Cell Reports 40.

77. Doherty, T. A., Baum, R., Newbury, R. O., Yang, T., Dohil, R., Aquino, M., Doshi, A., Walford, H. H., Kurten, R. C., Broide, D. H., et al. (2015). Group 2 innate lymphocytes (ILC2) are enriched in active eosinophilic esophagitis. Journal of Allergy and Clinical Immunology 136, 792–794.

78. Morgan, D. M., Ruiter, B., Smith, N. P., Tu, A. A., Monian, B., Stone, B. E., Virk-Hundal, N., Yuan, Q., Shreffler, W. G., and Love, J. C. (2021). Clonally expanded, GPR15-expressing pathogenic effector TH2 cells are associated with eosinophilic esophagitis. Science Immunology 6, eabi5586. 10.1126/sciimmunol.abi5586.

79. Yang, Y., Tresp, V., Wunderle, M., and Fasching, P. A. (2018). “Explaining Therapy Predictions with Layer-Wise Relevance Propagation in Neural Networks”. ICHI, 152–162. 10.1109/ICHI.2018.00025.

80. Minh, D., Wang, H. X., Li, Y. F., and Nguyen, T. N. (2022). Explainable artificial intelligence: a comprehensive review. Artificial Intelligence Review, 3503–3568.

81. Chen, C., Li, O., Tao, D., Barnett, A., Rudin, C., and Su, J. K. (2019). This looks like that: deep learning for interpretable image recognition. Advances in Neural Information Processing Systems 32.

82. Ali, S., Abuhmed, T., El-Sappagh, S., Muhammad, K., Alonso-Moral, J. M., Confalonieri, R., Guidotti, R., Del Ser, J., Díaz-Rodríguez, N., and Herrera, F. (2023). Explainable Artificial Intelligence (XAI): What we know and what is left to attain Trustworthy Artificial Intelligence. Information Fusion 99, 101805.

83. Buenrostro, J. D., Wu, B., Litzenburger, U. M., Ruff, D., Gonzales, M. L., Snyder, M. P., Chang, H. Y., and Greenleaf, W. J. (2015). Single-cell chromatin accessibility reveals principles of regulatory variation. Nature 523, 486–490.

84. Svensson, V. (2020). Droplet scRNA-seq is not zero-inflated. Nature Biotechnology 38, 147–150.

85. Maan, H., Zhang, L., Yu, C., Geuenich, M. J., Campbell, K. R., and Wang, B. (2024). Characterizing the impacts of dataset imbalance on single-cell data integration. Nature Biotechnology, 1899–1908.

86. Ioffe, S. and Szegedy, C. (2015). “Batch normalization: Accelerating deep network training by reducing internal covariate shift”. International conference on machine learning. pmlr, 448–456.

87. Nair, V. and Hinton, G. E. (2010). “Rectified linear units improve restricted boltzmann machines”. Proceedings of the 27th International Conference on Machine Learning (ICML-10), 807–814.

88. Loshchilov, I. and Hutter, F. (2019). “Decoupled weight decay regularization”. International Conference on Learning Representations.

89. Guo, C., Pleiss, G., Sun, Y., and Weinberger, K. Q. (2017). “On calibration of modern neural networks”. International conference on machine learning. PMLR, 1321–1330.

90. Pedregosa, F., Varoquaux, G., Gramfort, A., Michel, V., Thirion, B., Grisel, O., Blondel, M., Prettenhofer, P., Weiss, R., Dubourg, V., et al. (2011). Scikit-learn: Machine Learning in Python. Journal of Machine Learning Research 12, 2825–2830.

91. Luecken, M. D., Büttner, M., Chaichoompu, K., Danese, A., Interlandi, M., Müller, M. F., Strobl, D. C., Zappia, L., Dugas, M., Colomé-Tatché, M., et al. (2022). Benchmarking atlas-level data integration in single-cell genomics. Nature methods 19, 41–50.

92. Wolf, F. A., Angerer, P., and Theis, F. J. (2018). SCANPY: large-scale single-cell gene expression data analysis. Genome Biology 19, 15.

93. Ashburner, M., Ball, C. A., Blake, J. A., Botstein, D., Butler, H., Cherry, J. M., Davis, P., Dolinski, K., Dwight, S. S., Eppig, J. T., et al. (2000). Gene ontology: tool for the unification of biology. Nature genetics 25, 25–29.

94. Consortium, T. G. O. (2025). The Gene Ontology knowledgebase in 2026. Nucleic Acids Research 54, D1779–D1792. 10.1093/nar/gkaf1292. eprint: https://academic.oup.com/nar/article-pdf/54/D1/D1779/66009250/gkaf1292.pdf.

95. Benjamini, Y. and Hochberg, Y. (1995). Controlling the false discovery rate: a practical and powerful approach to multiple testing. Journal of the Royal statistical society: series B (Methodological) 57, 289–300.

96. Gayoso, A., Lopez, R., Xing, G., Boyeau, P., Valiollah Pour Amiri, V., Hong, J., Wu, K., Jayasuriya, M., Mehlman, E., Langevin, M., et al. (2022). A Python library for probabilistic analysis of single-cell omics data. Nature Biotechnology 40, 163–166.

